# The peripheral T cell population is associated with pneumonia severity in cynomolgus monkeys experimentally infected with severe acute respiratory syndrome coronavirus 2

**DOI:** 10.1101/2021.01.07.425698

**Authors:** Noriyo Nagata, Naoko Iwata-Yoshikawa, Kaori Sano, Akira Ainai, Nozomi Shiwa, Masayuki Shirakura, Noriko Kishida, Tomoko Arita, Yasushi Suzuki, Toshihiko Harada, Yasuhiro Kawai, Yasushi Ami, Shun Iida, Harutaka Katano, Seiichiro Fujisaki, Tsuyoshi Sekizuka, Hiroyuki Shimizu, Tadaki Suzuki, Hideki Hasegawa

## Abstract

The coronavirus disease 2019 (COVID-19), caused by severe acute respiratory syndrome coronavirus 2 (SARS-CoV-2), is a global pandemic that began in December 2019. Lymphopenia is a common feature in severe cases of COVID-19; however, the role of T cell responses during infection is unclear. Here, we inoculated six cynomolgus monkeys, divided into two groups according to the CD3+ T cell population in peripheral blood, with two clinical isolates of SARS-CoV-2: one of East Asian lineage and one of European lineage. After initial infection with the isolate of East Asian lineage, all three monkeys in the CD3+ low group showed clinical symptoms, including loss of appetite, lethargy, and transient severe anemia with/without short-term fever, within 14 days post-infection (p.i.). By contrast, all three monkeys in the CD3+ high group showed mild clinical symptoms such as mild fever and loss of appetite within 4 days p.i. and then recovered. After a second inoculation with the isolate of European lineage, three of four animals in both groups showed mild clinical symptoms but recovered quickly. Hematological, immunological, and serological tests suggested that the CD3+ high and low groups mounted different immune responses during the initial and second infection stages. In both groups, anti-viral and innate immune responses were activated during the early phase of infection and re-infection. However, in the CD3+ low group, inflammatory responses, such as increased production of monocytes and neutrophils, were stronger than those in the CD3+ high group, leading to more severe immunopathology and failure to eliminate the virus. Taken together, the data suggest that the peripheral T lymphocyte population is associated with pneumonia severity in cynomolgus monkeys experimentally infected with SARS-CoV-2.

**Author summary:** SARS-CoV-2 infection causes an illness with clinical manifestations that vary from asymptomatic or mild to severe; examples include severe pneumonia and acute respiratory distress syndrome. Lymphopenia, which is common in severe COVID-19 cases, is characterized by markedly reduced numbers of CD4+ T cells, CD8+ T cells, B cells, and natural killer cells. Here, we showed that cynomolgus monkeys selected according to the T cell populations in peripheral blood have different outcomes after experimental infection with SARS-CoV-2. These findings will increase our understanding of disease pathogenesis and may facilitate the development of animal models for vaccine evaluation.

## Introduction

Coronavirus disease 2019 (COVID-19), caused by a novel human coronavirus called severe acute respiratory syndrome coronavirus 2 (SARS-CoV-2), is a global pandemic that began in December, 2019 after cases of an unknown upper respiratory tract infection were reported in Wuhan, Hubei Province, China [1-3]. The World Health Organization declared a global pandemic on March 11, 2020; since then, the number of confirmed cases and the number of deaths has increased rapidly, reaching over 1 million by the end of September 2020 [4].

SARS-CoV-2 causes an illness with clinical manifestations ranging from an asymptomatic or mild infection to a serious illness (i.e., severe pneumonia and acute respiratory distress syndrome) [3, 5, 6]. Pathological studies suggest that SARS-CoV-2 infection of the lower respiratory tract causes disease directly [7, 8]. In addition, high expression of pro-inflammatory cytokines, including IL-6 and IL-1β, in serum from patients with severe COVID-19 suggest that immunopathological damage caused by an over-exuberant host response might contribute to poor outcomes [3, 9, 10]; this is similar to other coronavirus infections such as SARS and Middle East respiratory syndrome (MERS) [11-15]. Lymphopenia is a common characteristic of severe COVID-19; severe cases show a marked reduction in the numbers of CD4+ T cells, CD8+ T cells, B cells, and natural killer (NK) cells [3, 9, 10]. Because T cells may mediate early innate immune responses to virus infection [16], lymphopenia might be associated with severe disease. However, the role of T cell responses during COVID-19 infection is unclear.

Several experimental models, including cats, chickens, dogs, ducks, ferrets, mice, hamsters, macaque monkeys, and pigs, have been used to study COVID-19 [17, 18]. Cats, ferrets, human ACE2 transgenic mice, hamsters, and monkeys are all susceptible to SARS-CoV-2 after respiratory inoculation and all exhibit virus excretion from the upper respiratory tract and/or intestine [19-26]. These animals develop acute pulmonary lesions after inoculation with a high dose of virus, but clinical symptoms are mild. As in human cases of SARS, advanced age correlates with adverse outcomes in mice and macaque monkeys [27, 28]. However, cynomolgus monkeys do not show age-dependent differences in severity after experimental infection with SARS-CoV-2 [20].

Previously, we found that experimental infection of cynomolgus monkeys with human viral pathogens resulted in a few severe cases [29]. Pathophysiological analysis suggested that low populations of lymphocytes were related to the severe clinical symptoms after experimental infection with virus. Thus, we speculated that low T cell populations in peripheral blood might cause poor outcomes after SARS-CoV-2 infection of monkeys. To test this hypothesis, we selected monkeys according to the T cell population in peripheral blood, and infected them with an isolate of SARS-CoV-2 obtained from an individual who returned from Wuhan at the end of January 2020. We then monitored the clinical symptoms, immune responses, and lung pathology. In addition, we examined the effect of a previous infection with SARS-CoV-2 by reinfecting monkeys with a heterologous strain to evaluate whether it enhanced the symptoms of respiratory disease. We did this by re-challenging monkeys with another isolate, a “S-G614 variant strain”, isolated from a returnee from Europe at the end of March 2020. The results suggest that the peripheral T lymphocyte population in peripheral blood is related to severity of pneumonia in cynomolgus monkeys experimentally infected with SARS-CoV-2.

## Results

### Experimental infection of cynomolgus monkeys with SARS-CoV-2

An overview of the study design is shown in Figure 1A. Twenty-five female monkeys were used. Body weight was measured (S1A Fig) and blood samples obtained for use in a SARS-CoV-2 neutralization assay. All animals except one had undetectable (<1:4) levels of neutralizing antibodies; the exception had a titer of 1:4. The blood samples were also used to investigate the number of lymphocytes and the population of CD3+ cells within the total lymphocyte population (S1B Fig). After assigning animals into “CD3+ high” and “CD3+ low” groups, six cynomolgus monkeys were selected according to body weight (only monkeys weighing 3.5 kg or less were appropriate due to facility restrictions) and CD3+ cell count, and then used in the infection experiments.

**Fig. 1.**
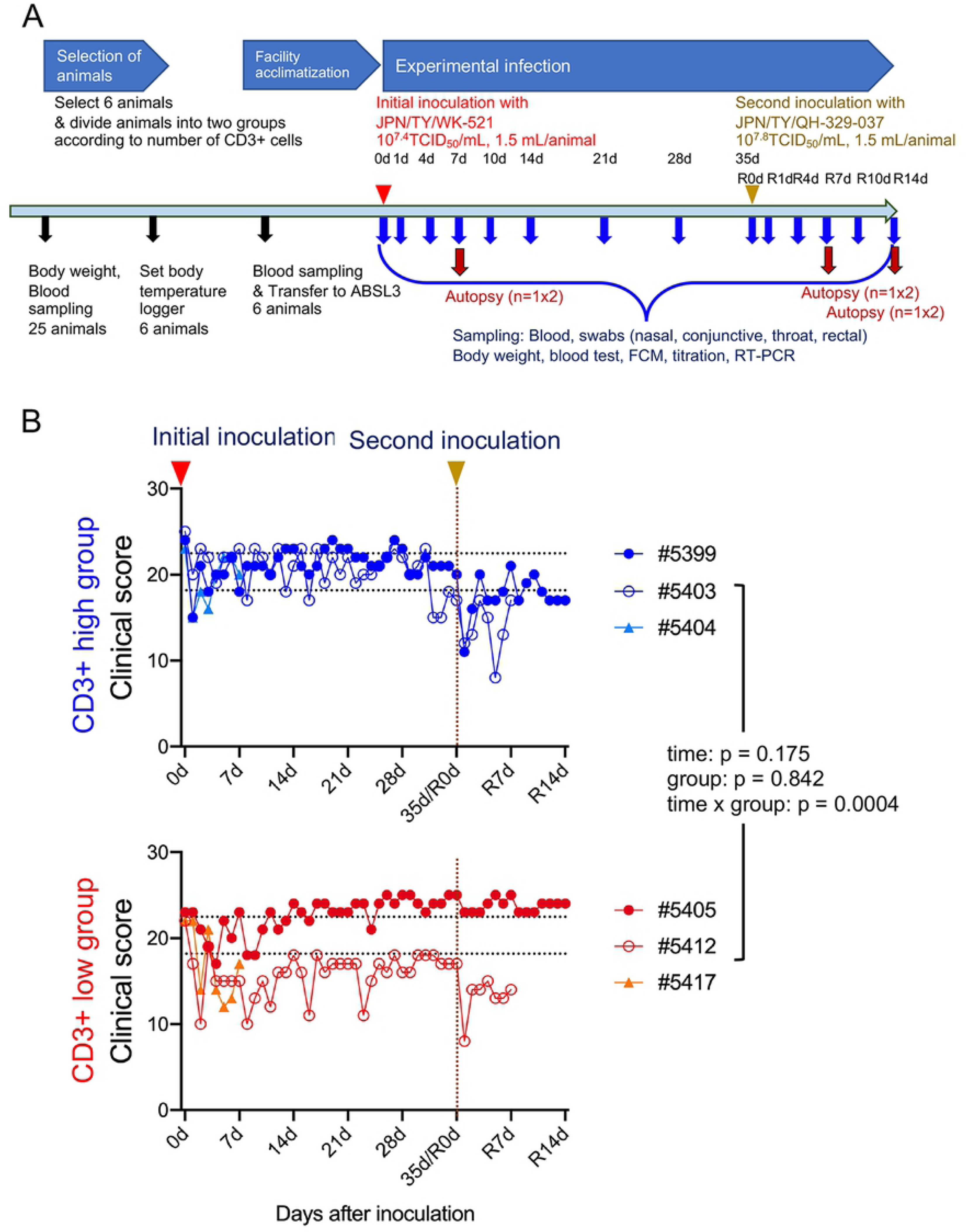
Study design and clinical scores in cynomolgus monkeys after inoculation of SARS-CoV-2. Study outline (A). Black arrows indicate preparation for experimental infection. Six 5-year-old monkeys were selected from 25 monkeys. Red and yellow arrow heads indicate virus inoculation. After assigning animals to “CD3+ high” and “CD3+ low” groups, six cynomolgus monkeys were infected with an isolate from East Asia (WK-521 strain) via a combination of intranasal (0.125 mL, sprayed into the right nostril), conjunctival (0.1 mL, dropped into the right eye), and intratracheal (1.275 mL virus solution plus 2 mL saline via a catheter) inoculation. After the initial inoculation, body weight was measured, and samples were collected at various time points (blue arrows). Red arrows denote autopsy at 7 or 14 days after the first or second inoculation (n = 1 per group at each time point). Four monkeys received a second inoculation with an isolate from Europe (QH-329-037 strain). (B) Clinical scores of cynomolgus monkeys inoculated with SARS-CoV-2. Cool (blue and aqua) and warm (red and orange) colored symbols and lines indicate data from the CD3+ high group and CD3+ low group animals, respectively. After transfer to the ABSL3 facility, the monkeys were observed once daily for clinical signs and scored accordingly. Black dashed lines on the horizontal axis indicate the range of clinical scores recorded during the ABSL3 facility acclimatization period. Each dot/line represents data from an individual animal after initial inoculation with SARS-CoV2. The brown dashed line on the vertical axis indicates the day of the second inoculation.

In this study, we used two isolates of SARS-CoV-2 from Japan: one of East Asian lineage obtained at the end of January 2020, and another of European lineage obtained at end of March 2020 (Table 1). The first inoculation with the isolate of East Asian lineage (2019-nCoV/Japan/TY/WK-521/2020, referred to as WK-521) was administered via a combination of the intranasal (0.25 mL, spray into right nostril), conjunctival (0.1 mL, drop on right eye), and intratracheal (1 mL virus solution plus 2 mL saline using a catheter) routes under ketamine-xylazine anesthesia; the monkeys were observed once daily for clinical signs and scored using a clinical scoring system (dietary intake, including pellets and fruits, drinking, attitude in front of regular observers, and stool consistency: the total score was the sum of all five component scores (i.e., 0–5 × 5). (Fig. 1B).

**Table 1.** Clinical isolates of SARS-CoV-2 used in this study.

| Strain | Origin |  | Accession no. | GISAID Clade* /<br>region of exposure | Passage history for the animal experiment in this study |  |  |
| --- | --- | --- | --- | --- | --- | --- | --- |
|  | Collection date | Specimen |  |  | Cell | Propagation | Mycoplasma*** |
| 2019-nCoV/Japan /TY<br>/WK-521/2020 | Jan 31, 2020<br>Returnee from Wuhan | Throat swab | EPI_ISL_408667 | S / East Asia | VeroE6<br>/TMPRSS2 | 8 passages** | Negative |
| hCoV-19/Japan/<br>QH-329-037/2020 | 29 Mar 2020<br>Returnee from EU | Throat swab | EPI_ISL_529135 | G / Europe | VeroE6<br>/TMPRSS2 | 2 passages | Positive |

GSAID, Global Initiative on Sharing All Influenza Data.
\*GSAID clade referred from "Clade and lineage nomenclature, July 4, 2020"
https://www.gisaid.org/references/statements-clarifications/clade-and-lineage-nomenclature-aids-in-genomic-epidemiology-of-active-hcov-19-viruses/.
\*\*WK-521 was isolated using VeroE6/TMPRSS2 cells unexpectedly contaminated with *Mycoplasma hyorhinis* and *Mycoplasma arginini* (Matsuyama et al., 2020). Anti-mycoplasma reagents (MC-210, 0.5 µg/mL; Waken, Kyoto, Japan) were used to eradicate mycoplasma contamination from the cells and virus stock during virus propagation from passages 4 to 6.
\*\*\*Mycoplasma contamination was detected by PCR (TaKaRa PCR Mycoplasma Detection Set, Takara, Shiga, Japan).

All monkeys showed reduced appetite, drank less, and became more lethargic within 4–10 days after the initial inoculation. Two animals (#5412 and #5417) from the CD3+ low group showed lower clinical scores than that for the CD3+ high group from 5 to 14 days p.i. Monkey #5412 became lethargic around 10 days after the initial inoculation but ate a piece of fruit every day; therefore, we decided not to euthanize this animal. Indeed, the monkey recovered from severe illness at around 14 days p.i. After the second inoculation with another isolate of European lineage (hCoV-19/Japan/ QH-329-037/2020, referred to as QH-329-037) at 35 days after the initial inoculation (referred to as R0d in Fig. 1), all monkeys except #5405 showed a reduced clinical score and recovered within 1 week. No obvious body weight loss was observed; indeed, monkey #5405 gained weight (S2A Fig). In all monkeys, body temperature spiked 1 day after the initial inoculation but then returned to normal (the exception was monkey #5405) (S3 Fig). Monkey #5405 continued to have a slightly higher temperature than before the initial inoculation. At 1 day after the second inoculation, two monkeys from the CD3+ high group (#5399 and #5403) showed a spike in body temperature. Biochemical markers (globulin: Glob, albumin: ALB, and glucose) suggested changes in nutritional status after both the initial and second inoculations (S2B Fig).

Two monkeys from the CD3+ low group showed low hemoglobin (HGB) levels: one at 7 days (monkey #5417, at the time of planned autopsy) and one at 10 days (monkey #5412) after initial inoculation (Fig. 2A). Red blood cell (RBC) counts and hematocrit levels were also low in these monkeys (S4A Fig).

**Fig. 2.**
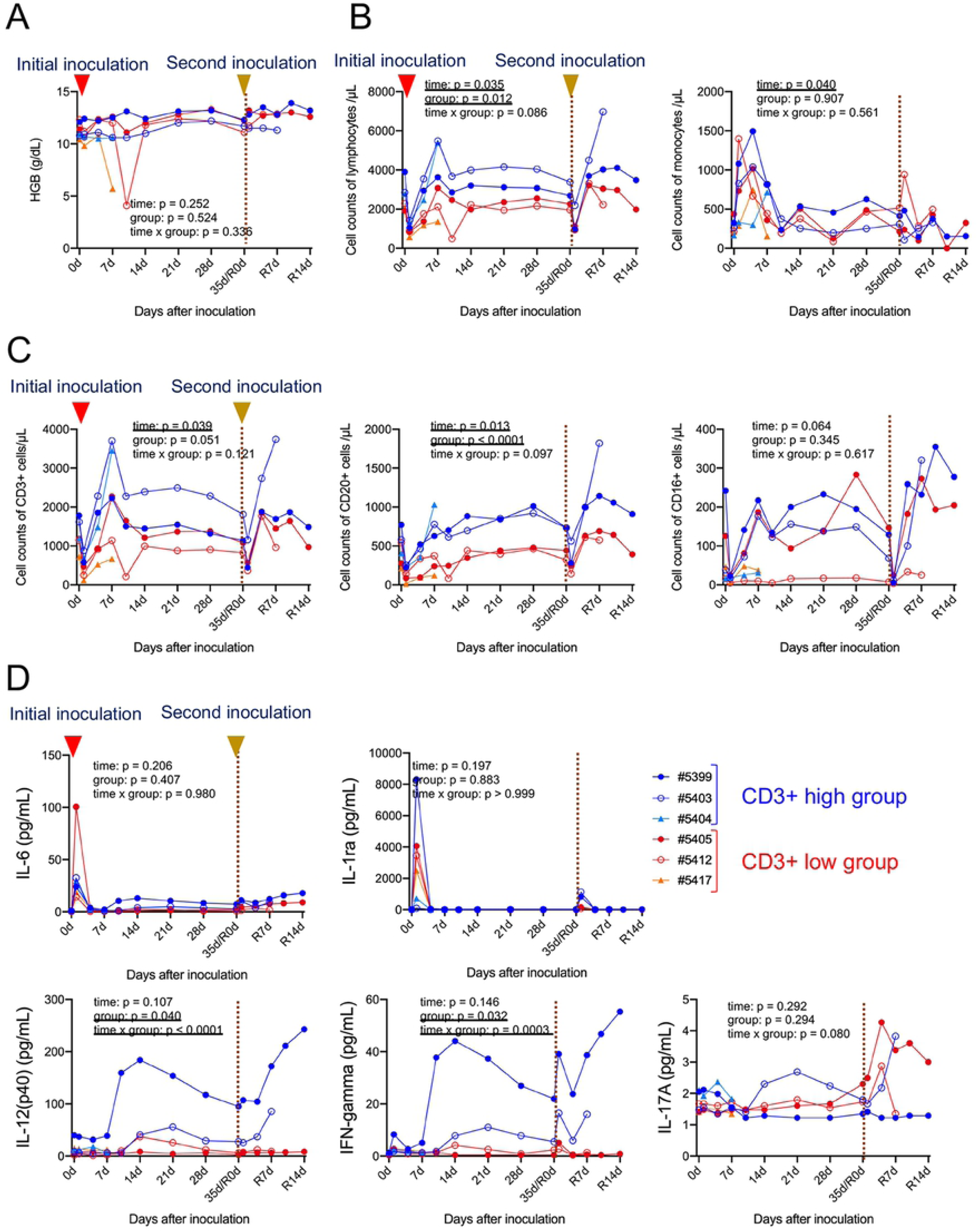
Hematological examination of cynomolgus monkeys inoculated with SARS-CoV-2. Hemoglobin (HGB) in EDTA-treated whole blood samples was examined at various time points after inoculation (A). Absolute numbers of lymphocytes and monocytes in EDTA-treated whole blood samples were determined at various time points after inoculation (B). Leukocyte differentiation (e.g., CD3, CD20, and CD16) at various time points after inoculation was examined by flow cytometry (C). Cytokine and chemokine levels in serum from each cynomolgus monkey inoculated with SARS-CoV-2 (D). Representative cytokines were profiled by multiplex analysis. Assays were performed using unicate samples at each time point. Cool (blue and aqua) and warm (red and orange) colored symbols and lines indicate data from the CD3+ high and CD3+ low groups, respectively. Each dot/line represents data from an individual animal. The brown dashed line on the vertical axis indicates the day of the second inoculation.

### Immune responses in cynomolgus monkeys inoculated with SARS-CoV-2

All monkeys showed transient lymphopenia at 1 day p.i., after which lymphocyte counts increased within the next 7 days (Fig. 2B). After the second inoculation, lymphocyte counts in all monkeys decreased at Day 1 p.i. before recovering again. Mixed-effects models for repeated measures analysis revealed significant differences in the number of lymphocytes between the two groups. By contrast, monocyte counts after the first injection increased before falling again within 7 days p.i. (Fig. 2B). After the second inoculation, monocyte counts did not change significantly. Various changes in the numbers of other leukocytes, including neutrophils, eosinophils, basophils, were seen during infection (S4B Fig).

Flow cytometry analysis revealed that changes in the overall lymphocyte count were due to changes in the number of CD3+ T cells (Fig. 2C). In both groups, CD20+ B cell counts dropped at 1 day p.i. and then increased gradually until 28 days p.i., but interestingly, there was a significant difference between CD20+ B cell counts in the CD3+ high and low groups (Fig. 2C). After the second inoculation, the number of CD20+ B cells in both groups fell, before increasing again. Three monkeys showed high CD16+ NK cell counts at 4 days after the initial inoculation (Fig. 2C). After the second inoculation, CD16+ NK cells numbers in all monkeys were higher than after the initial inoculation, although numbers remained low in monkey #5412. There was a significant difference in the number of CD3+CD4+ T cells between the two groups (S4C Fig). CD3+ cells, including CD4+ and CD8+ T cell counts, peaked at 7 days after the initial inoculation, but cell numbers increased rapidly after the second inoculation.

Levels of IL-6, interleukin 1 receptor antagonist (IL-1ra), monocyte chemotactic protein-1 (MCP-1), IL-15, IL-2, and macrophage inflammatory protein-1 beta (MIP-1β) in serum peaked at 1 day after the initial inoculation; levels also increased at 1 day after the second inoculation, although the increase was smaller in both groups (Fig. 2D and S5A Fig). Levels of helper T cell (Th cell)-related cytokines, such as IL-12/23 (p40), interferon gamma (IFN-γ), tumor necrosis factor alpha (TNF-α), IL-13, IL-10, and IL-17 increased from Day 10 post-initial inoculation, peaking at Day 14 or 21; expression increased rapidly (within 7 days) after the second inoculation (Fig. 2D and S5B Fig). The kinetics of Th cell-related cytokine responses (except IL-17) were faster in the CD3+ high group than in the CD3+ low group. Dynamic changes in transforming growth factor alpha (TGF-α) and IL-8 levels were also observed in both groups during infection (S5C Fig).

### Virus shedding by cynomolgus monkeys inoculated with SARS-CoV-2

After the initial inoculation with isolate WK-521, clinical samples (conjunctiva, nasal, throat, and rectal swabs) were collected. Viral RNA was detected by real-time RT-PCR, and infectious virus was detected by culture with TMPRSS2-Vero E6 cells (Fig. 3). The result revealed that two monkeys excreted infectious virus from the upper respiratory tract (nasal and throat swabs from #5404) or intestine (rectal swab from #5412) after the initial inoculation. Real-time RT-PCR confirmed viral replication in the upper respiratory tract and intestine by detecting viral subgenomic mRNAs in swab samples that were positive for viral RNA [30]. Actively-infected cells were detected in nasal swabs from two monkeys (#5403 and #5404) and in a rectal swab from one monkey (#5412) (Fig. 3 and S6 Fig). After the second inoculation, none of the monkeys excreted infectious virus, although viral subgenomic mRNA was detected in nasal (#5403) and rectal (#5412) swabs (Fig. 3).

**Fig. 3.**
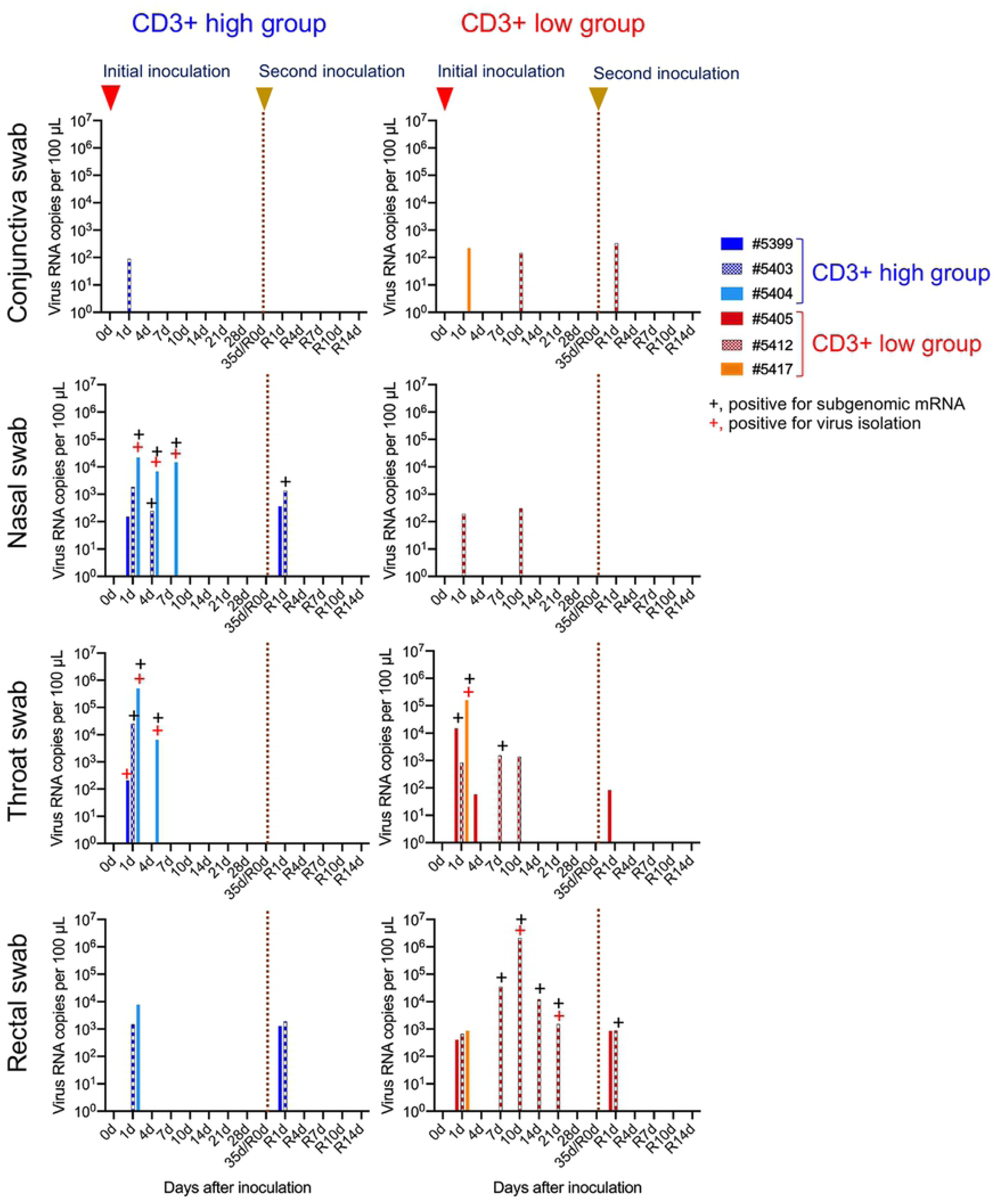
Detection of virus excretion in clinical samples from cynomolgus monkeys inoculated with SARS-CoV-2. Six cynomolgus monkeys were used in this study. Cool (blue and aqua) and warm (red and orange) colored bars indicate data from CD3+ high and CD3+ low groups, respectively. Each bar represents data from an individual animal. After initial viral inoculation with the WK-521 strain, clinical samples (conjunctiva, nasal, throat, and rectal swabs) were collected. The second inoculation with QH-329-037 strain was performed 35 days after the first inoculation. + indicates samples that were positive for subgenomic mRNA (black) or virus (red). The brown dashed line on the vertical axis indicates the day of the second inoculation.

### Seroconversion after experimental infection with SARS-CoV-2

No monkeys, including #5404 and #5417 euthanized on Day 7 p.i., showed seroconversion within 7 days p.i. Neutralizing antibodies were detected from 10 days (monkey #5399 in the CD3+ high group) or 14 days (the other three monkeys in both groups) after the initial inoculation, peaking at 21 days p.i. in the CD3+ high group and 28 days p.i. in the CD3+ low group (Fig. 4A). Within 35 days p.i. (35d/R0d in Fig. 4A), the neutralizing antibody titer in monkeys #5399 and #5403 from the CD3+ high group fell slightly; overall, the antibody titers were higher in the CD3+ low group than in the CD3+ high group. After the second inoculation, neutralizing antibody titers increased rapidly at 4 days (R4d) p.i., peaking at 1:640 at 7 days (R7d) p.i., in all monkeys from both groups. Monkeys #5403 and #5412 were euthanized at R7 days p.i. After this time point, the titers in monkeys #5399 and #5405 fell slightly to 1:320 at 14 days. Sidak’s multiple comparisons test after application of a mixed-effects models for repeated measures analysis revealed a significant difference in neutralizing antibody titers between the two groups. Serum obtained from the monkeys showed cross-reactivity with both strains of virus (S1 Table).

**Fig. 4.**
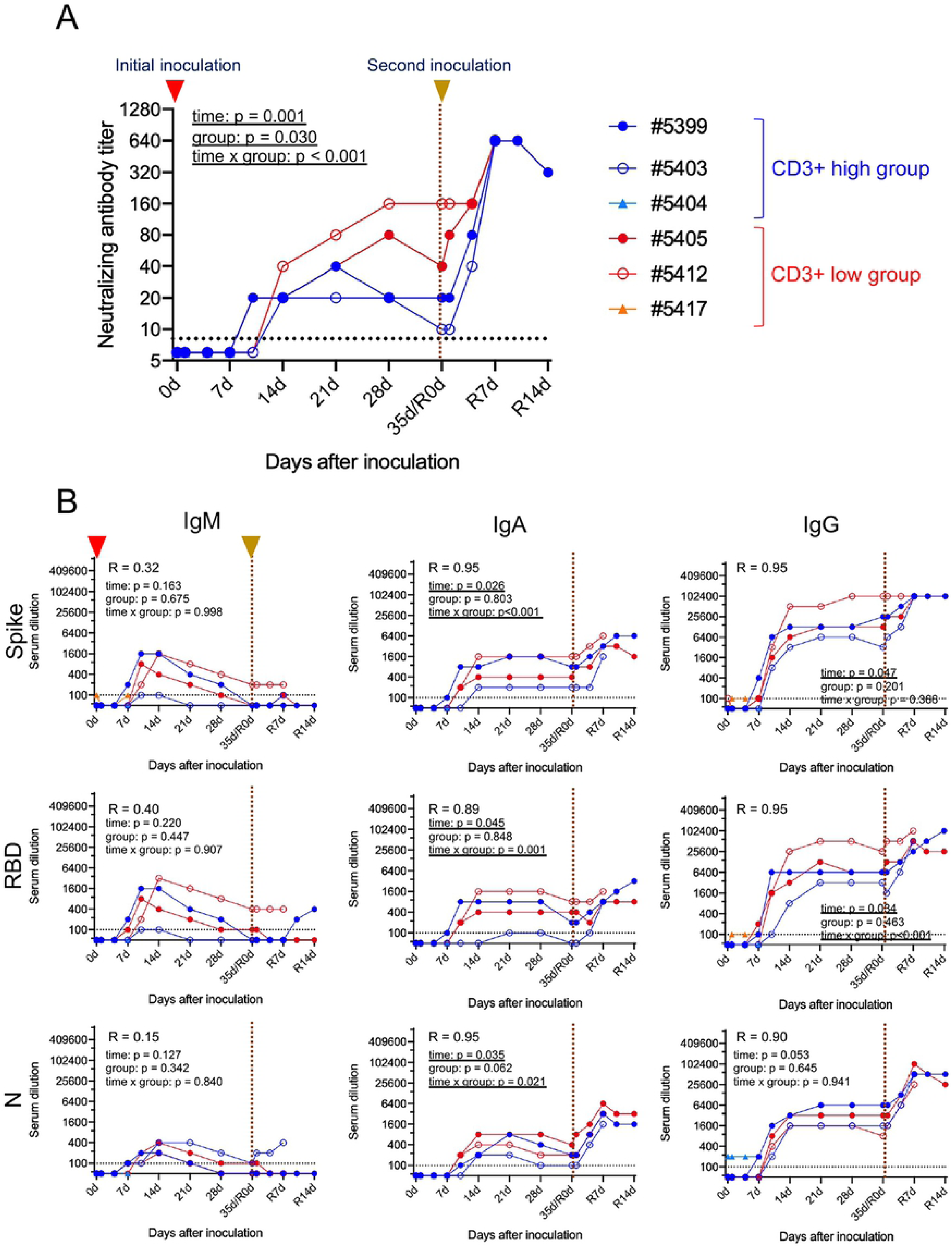
Seroconversion after SARS-CoV-2 inoculation. Neutralizing antibody titers (against the WK-521 strain) in sera (A). Antibody subclasses and specificity for the spike (S), receptor binding domain (RBD), and nucleocapsid (N) proteins were assessed using in-house IgM, IgA, and IgG ELISAs (B). Cool (blue and aqua) and warm (red and orange) colored symbols and lines indicate data from the CD3+ high and CD3+ low groups, respectively. Each dot/line represents data from an individual animal. R, correlation coefficient (Spearman’s correlation analysis) between the neutralization and ELISA tests. The brown dashed line on the vertical axis indicates the day of the second inoculation.

We also used in-house IgM, IgA, and IgG enzyme-linked immunosorbent assay (ELISAs) to examine antibody isotypes and their binding to the spike (S), receptor binding domain (RBD), and nucleocapsid (N) proteins (Fig. 4B). At 7 or 10 days p.i., S-, RBD-, and N protein-specific IgM, IgA, and IgG antibody titers increased in both groups. Spearman’s correlation analysis revealed that the IgA and IgG responses correlated with the neutralizing antibody response (R > 0.8). High levels of IgG antibodies specific for the S and RBD proteins were observed in monkey #5412, which showed prolonged excretion of infectious virus from the intestine after the initial inoculation.

### Transcriptomic analyses of peripheral whole blood from monkeys inoculated with SARS-CoV-2

Transcriptomic analyses were conducted using RNA extracted from peripheral whole blood samples collected at different time points: before initial inoculation (Day 0), after initial inoculation (Days 1, 4, and 7), before the second inoculation (R0), and after the second inoculation (R1, R4, and R7). Gene expression was compared between samples collected from animals before (Day 0) and after (Days 1, 4, 7, R0, R1, R4, and R7) virus inoculation to identify differentially expressed genes (S7A Fig). The results revealed that 331 genes were upregulated significantly, while 176 genes were downregulated significantly, after virus infection. Among the 507 differentially expressed genes, 78 were related to the immune response (S7B Fig). Next, we conducted gene set enrichment analyses using samples collected from the CD3+ high and CD3+ low groups after (Days 1, 4, 7, R0, R1, R4, and R7) virus infection (Fig. 5A). Expression of genes encoding neutrophil-, monocyte-, and inflammatory signal-related modules were downregulated to a greater extent in the CD3+ high group than in the CD3+ low group (green dots in Fig. 5A), whereas expression of genes encoding B cell-related modules was upregulated to a greater extent in the CD3+ high group than in the CD3+ low group (gray dots in Fig. 5A). Furthermore, to evaluate differences in transcriptomic profiles over time, we conducted gene set enrichment analyses at baseline (before virus infection, Day 0) and at different time points after virus infection (Days 1, 4, 7, R0, R1, R4, and R7) (Fig. 5B). The results revealed significant upregulation of genes encoding innate anti-viral immune system-related modules (yellow dots in Fig. 5B) on Days 1 and 4 in both the CD3+ high and CD3+ low groups. Of note, upregulation of genes encoding inflammation-related modules (green & red dots in Fig. 5B) and downregulation of genes encoding T cell-related modules (black dots in Fig. 5B) were more prominent in the CD3+ low group than in the CD3+ high group. Upregulation of innate immune response-related genes was observed following re-infection with virus, although no alteration in expression of T cell- and B cell-related genes was observed. Overall, expression of more immune response-related modules was altered significantly in the CD3+ low group compared with the CD3+ high group, suggesting a difference in the magnitude of the immune response between the two groups following virus infection.

**Fig. 5.**
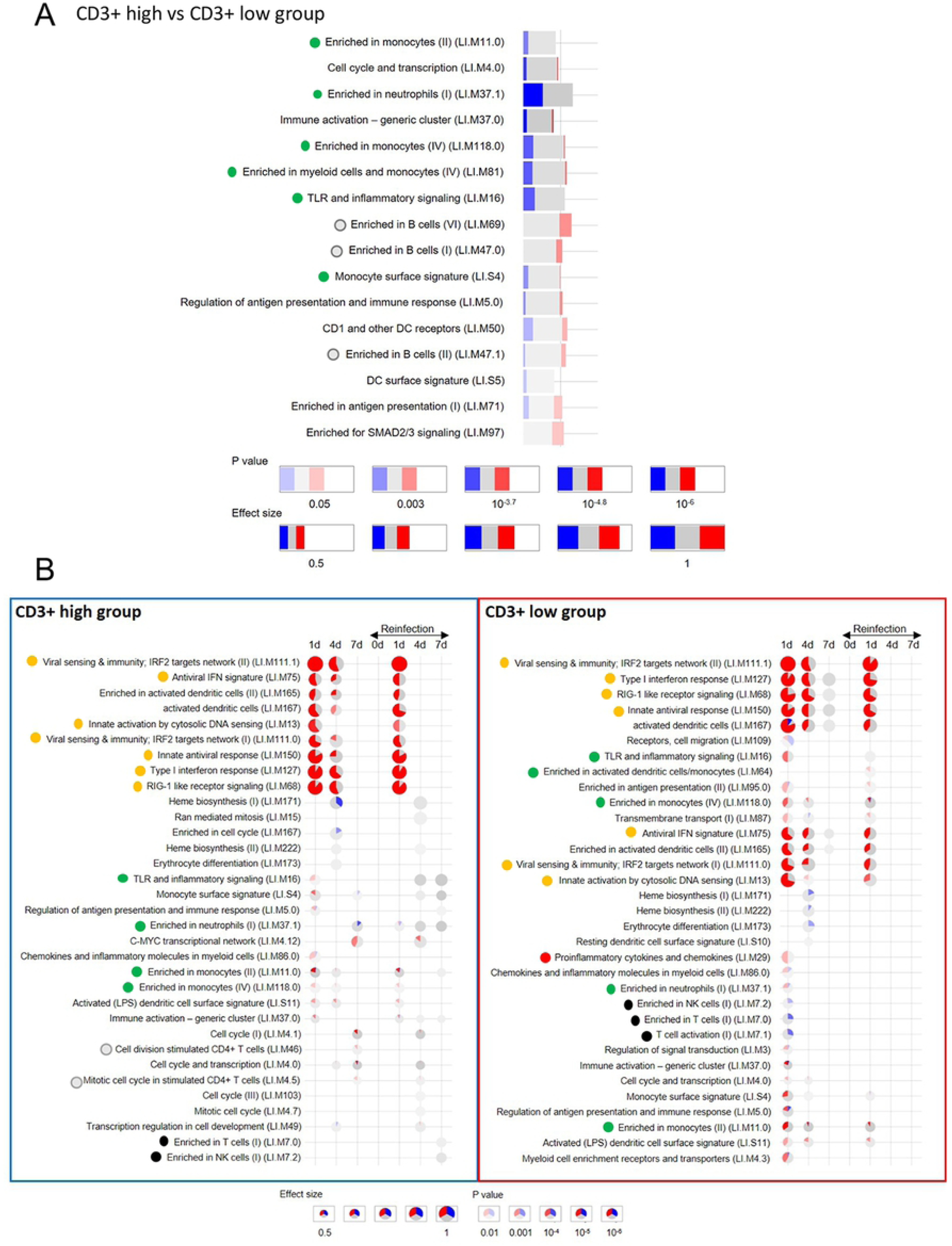
Transcriptome analysis of blood samples obtained after SARS-CoV-2 inoculation. Gene set enrichment analysis was performed on samples from the CD3+ high group and CD3+ low group samples after (Days 1, 4, 7, R0, R1, R4, and R7) virus infection (A). The name of each significantly enriched module is listed, along with the module ID (in brackets) (*P* < 0.01). Green and gray dots indicate inflammation- and B cell response-related modules, respectively. Red and blue indicate the proportion of genes in a particular module that is upregulated or downregulated in the CD3+ high group compared with the CD3+ low group. Each module is represented by a box, where the width is proportional to the effect size (AUROC value calculated from the number of genes in the module and ranking by the Cerno test), while brighter colors indicate lower *P*-values. Gene set enrichment analysis in the CD3+ high group (left panel) and CD3+ low group (right panel) at different time points after virus infection (Days 1, 4, 7, R0, R1, R4, and R7) compared with baseline (before virus infection: Day 0) (B). The name of each significantly enriched module name is listed along with module ID (in brackets) (*P* < 0.01). Yellow, green & red, gray, and black dots indicate modules related to innate immunity, inflammation, CD4+ T cell response, and T & NK cell responses, respectively. Red and blue indicate the proportion of genes in a particular module that is upregulated or downregulated in the CD3+ high group compared with the CD3+ low group. Each module is represented as a pie chart, where the size is proportional to the effect size (AUROC value calculated from the number of genes in the module and ranking by the Cerno test), while brighter colors indicate lower *P-*values.

### Distribution of viral RNA in monkey tissues at the experimental end-point

At the experimental end-point, tissue samples were also collected to detect viral RNA, subgenomic mRNA, and infectious virus (Fig. 6 and S6B Fig). Two monkeys (#5404 and #5417) euthanized at 7 days after initial inoculation had viral RNA and/or subgenomic mRNA in the upper and lower lobe of the lungs. At 7 and 14 days after the second inoculation (R7d and R14d), two monkeys (#5412 and #5405) from the CD3+ low group had viral RNA and/or subgenomic mRNA in the lower lobe of the lungs and in the trachea. Monkey #5412 excreted the virus in rectal swabs (Fig. 3) and had detectable viral RNA and/or subgenomic mRNA in the large intestine and mesenteric lymph nodes at 7 days after the second inoculation. High levels of viral RNA were detected in the tonsil and subcarinal lymph nodes of monkeys in both the CD3+ high and low groups at various time points. No infectious virus was isolated from tissue samples using TMPRSS2/VeroE6 cells. Because it was difficult to distinguish cytopathic effects (CPE) from cytotoxicity caused by the tissue homogenate, we performed blind passage of TMPRSS2/VeroE6 cells in the presence of culture supernatant from the first inoculation plus 10% tissue homogenate; however, there were no distinct CPEs within 5 days p.i., after the second blind passage.

**Fig. 6.**
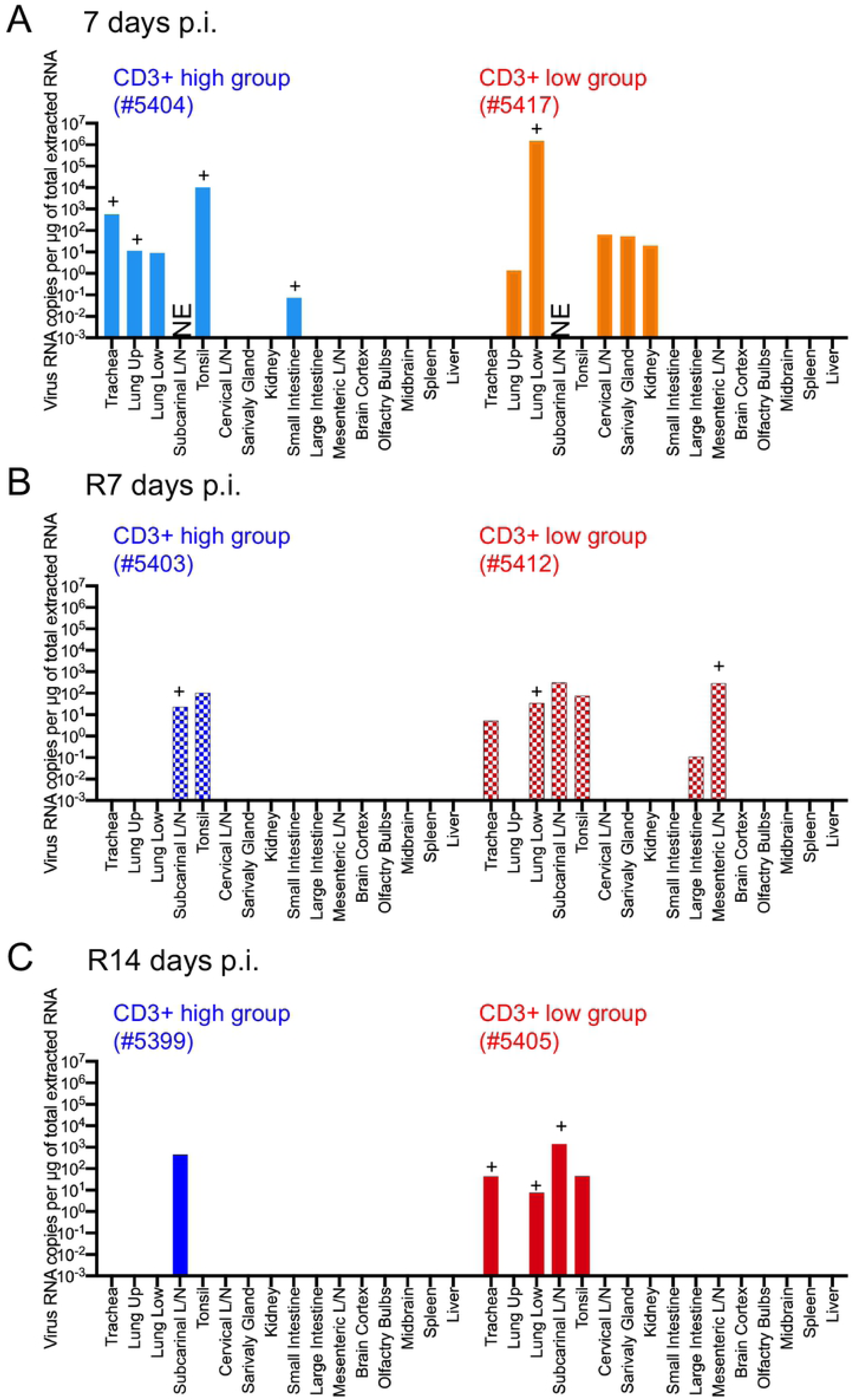
Detection of virus RNA in tissue samples from cynomolgus monkeys inoculated with SARS-CoV-2. Tissue samples were obtained from monkeys at 7 days post-inoculation with WK-521 strain (#5404 and #5417), and at 7 days (#5403 and #5412) or 14 days (#5399 and #5405) after re-infection with QH-329-037 strain. Cool (blue and aqua) and warm (red and orange) colored bars indicate data from the CD3+ high group and CD3+ low group, respectively. Each bar represents data from an individual animal.

As mentioned above, we used a heterologous strain of the virus for the second infection. To identify single nucleotide variations and the major population of virus in monkey tissues, we used the next generation sequencer MiSeq to obtain the entire length of the viral genome. Thirteen RNA samples obtained from tonsil, mesenteric lymph nodes, and lung tissues from infected monkeys were used for analysis; however, four samples did not meet the quality standards. The read number obtained for six samples of tonsil and lung returned only partial viral genome sequences (S2 Table). In the end, only three samples were suitable for genome sequencing; the sequences obtained from these samples were compared with the Wuhan-Hu-1 genome sequence (accession no. MN908947.3) as a reference (Table 2). The sequence data have been deposited in the DNA Data Bank of Japan (DDBJ) Sequence Read Archive, under submission ID DRA011219 (BioSample accessions: SAMD00261559 – 00261561). In addition, because the number of reads was sufficient at the D614G position (>200), a genetic population analysis of the D614G variant was performed in six samples (S3 Table).

**Table 2.** SARS-CoV-2 variants in tissue samples from monkeys after experimental infection.

| Group | Stain (Accession no.) | Nucleotide position, reference: Wuhan-Hu-1 (accession no. MN908947.3) |  |  |  |  |  |  |  |  |  |  |  |  |  |
| --- | --- | --- | --- | --- | --- | --- | --- | --- | --- | --- | --- | --- | --- | --- | --- |
|  |  | 1648 | 2662 | 4185 | 4456 | 5497 | 8782 | 11942 | 12334 | 13548 | 16596 | 18755 | 18804 | 21886 | 23403 |
| Region |  | ORF1a |  |  |  |  |  |  |  | ORF1b |  |  |  | S |  |
| Nonsynonymous mutation |  | - | - | G1307A | - | - | - | Q3893* | - | - | - | P1763L | - | - | D614G |
| 1st inoculum | WK-521 (EPI_ISL_408667) | C | T | G | C | C | T | C | A | C | C | C | C | T | A |
| CD3+ high<br>at 7 dpi | #5404_Tonsil<br>(SAMD00261560) | - | - | - | - | - | - | - | - | - | - | - | - | - | - |
| CD3+ low<br>at R7 dpi | #5412_Lung<br>(SAMD00261561) | T | - | - | T | T | T | - | del | T | T | T | T | - | - |
|  |  | 100%* |  |  | 100% | 62% | 60% |  | 100% | 100% | 100% | 51% | 50% |  |  |
|  | #5412_Mesenteric lymph node<br>(SAMD00261559) | - | - | C | - | - | - | T | - | - | - | - | - | C | - |
|  |  |  |  | 61%* |  |  |  | 55% |  |  |  |  |  | 100% |  |
| 2nd inoculum | QH-329-037 (EPI_ISL_529135) | - | C | - | - | - | C | - | - | - | - |  |  | - | G |

\*Percent nucleotide polymorphism; del, deletion.

The results revealed that the viral genome obtained from the tonsil of monkey #5404 after initial inoculation did not harbor any mutations (threshold = 50%). However, the viral genome isolated from the lung of monkey #5412 harbored nine single nucleotide polymorphisms (SNPs), including seven silent point mutations, one deletion resulting in a frameshift mutation in the ORF1ab region, and a nonsynonymous mutation in ORF1b. The most common base change was C > T. In addition, the genome isolated from mesenteric lymph nodes from monkey #5412 harbored three SNPs, including two nonsynonymous mutations in the ORF1a region and a synonymous mutation in the S region. The major sequence in these two isolates was derived from WK-521, suggesting that the original inoculum replicated and resided in the respiratory tract and intestine of monkey #5412, even after the second inoculation with the heterologous strain. The viral genome obtained from the tonsil of monkey #5403 after the second inoculation harbored a D614G mutation in the S region, suggesting the presence of QH-329-037 in the tonsil after the 2nd inoculum (S3 Table). The viral genome also obtained from the tonsil of #5405 after the second inoculation harbored a D614G mutation in the S region, suggesting the presence of QH-329-037 in the tonsil. Interestingly, the genome obtained from the subcarinal lymph node of monkey #5405 did not harbor the D614G mutation, suggesting that the original inoculum was maintained in the accessory lymph node of the lungs. These results suggest that the initially inoculated virus (WK-521) was maintained in the lungs and/or accessory lymph nodes, and that the second inoculated virus (QH-329-037) was eliminated from the lungs of these monkeys soon after the second inoculation.

### Pathology of SARS-CoV-2 infection in cynomolgus monkeys inoculated with SARS-CoV-2

Gross pathology of lungs from monkeys at each end-point is shown in Fig. 7A. Obvious gross lung lesions observed in monkey #5417 at 7 days after the initial inoculation (Fig. 7A, red and white arrows). After the second inoculation, enlargement of the subcarinal lymph nodes was seen in three monkeys, except #5405 (Fig. 7A, yellow arrows). Histopathological analysis revealed varying degrees of alveolar damage in monkeys #5404 and #5417 at 7 days after the initial inoculation (Fig. 7B). Lung tissue from monkey #5404 showed multifocal, slight to mild, interstitial pneumonia, with mononuclear cell aggregates in the alveoli (Fig. 7B, upper row). Monkey #5417 developed more severe interstitial pneumonia, with pulmonary edema comprising degenerated cells and polymorphonuclear leukocytes (Fig. 7B, lower row). Proliferating type II cells overlying the pulmonary walls were observed within the lesions. CD3+ lymphocytes and CD68+ macrophages were present in the alveoli. The lesions in the lungs of monkey #5417 contained predominantly CD68+ macrophages rather than CD3+ cells (Fig. 7C).

**Fig. 7.**
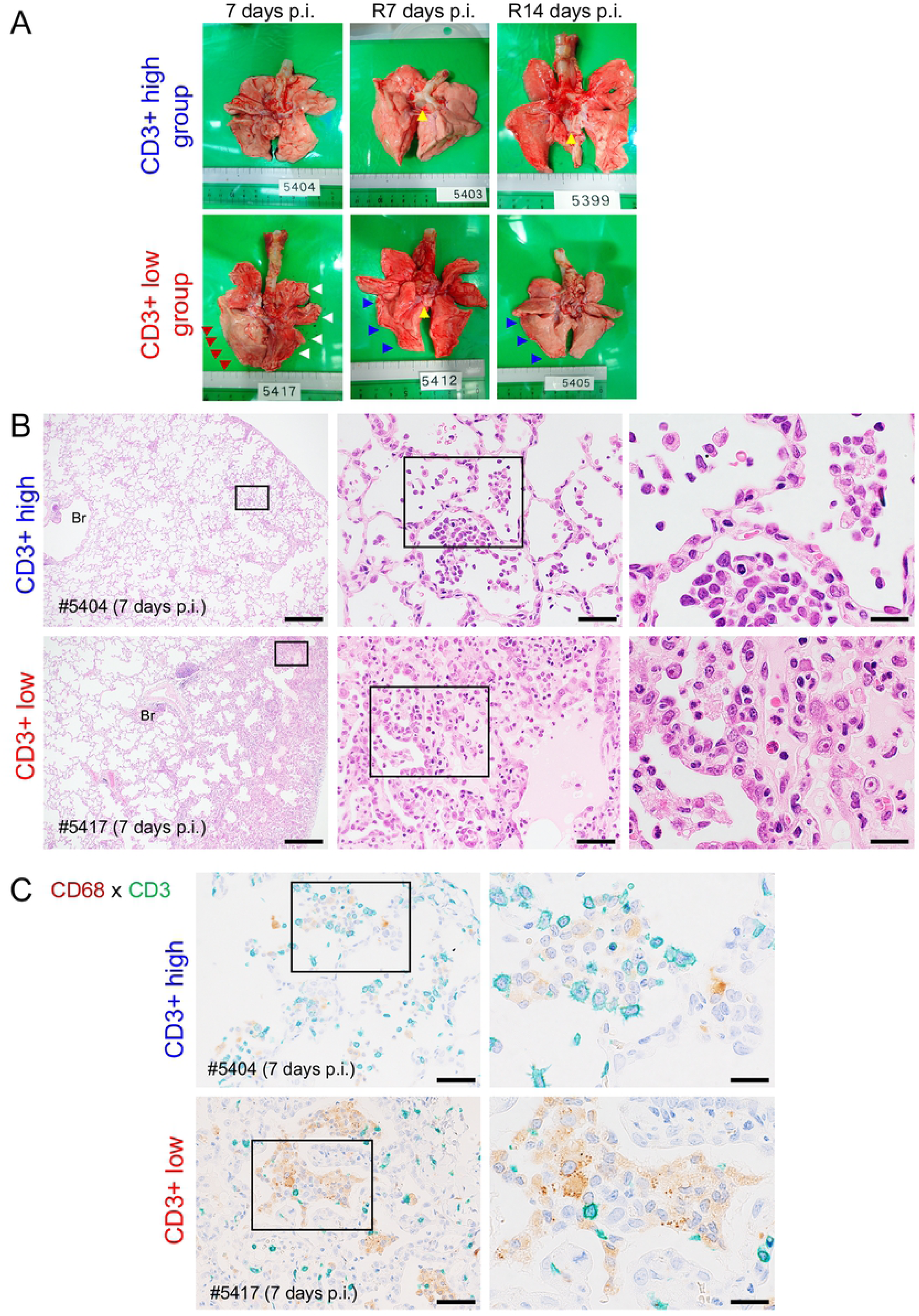
Pathology of cynomolgus monkeys inoculated with SARS-CoV-2. (A) Gross pathology of lungs from monkeys at 7 days post-inoculation with WK-521 strain (#5404 and #5417), and at 7 days (#5403 and #5412) or 14 days (#5399 and #5405) after re-infection with QH-329-037 strain. Ischemic changes and consolidation were observed in the lower lobe of the right lung of monkey #5417 (red arrows). Other lobes showed congestion and collapse (white arrows). Yellow arrows indicate swollen lung lymph nodes in monkeys #5403, #5412, and #5399. Atrophic changes are seen in the pulmonary margin in monkeys #5412 and #5405 (blue arrows). (B) Representative histopathology of lungs from monkeys at 7 days post-inoculation with WK-521 strain (#5404 and #5417). Collections of mononuclear cells were seen in the airspaces of the middle lobe of the right lung of monkey #5404 (B, upper row). Pulmonary edema with polymorphonuclear leukocyte infiltration and proliferating type II cells overlying pulmonary walls were observed in the lower lobe of the right lung of monkey #5417 (B, lower row). Scale bars: 500 µm (left column), 50 µm (middle column), and 20 µm (right column). Hematoxylin and eosin staining (H&E). (C) Double immunohistochemistry identified cell collections in alveolar air spaces at 7 days after the initial inoculation (upper row from #5404; lower row from #5417). Infiltrating cells were CD68+ (brown) or CD3+ (green). Bars in C, 50 µm (left) and 20 µm (right). An anti-CD68 rabbit polyclonal antibody (brown) and an anti-CD3-monoclonal antibody (green) were used for IHC in C.

Double immunohistochemistry revealed high expression of ACE2 on the surface of the pulmonary bronchi, but staining was very weak in the alveoli (S8 Fig, upper row); there was no merging of virus antigen (S8 Fig, brown) and ACE2 (S8 Fig, green) signals in either area. In addition, ACE2 was strongly expressed by hyperplastic type II pneumocytes in the pulmonary lesions (S8 Fig, lower row).

Monkeys euthanized after the second inoculation had slight focal interstitial inflammation, with macrophages and lymphocytes in the alveoli but no evidence of viral antigens (S9 Fig).

Supplementary figure 10 shows representative examples of histopathology of the lungs and extrapulmonary organs. Hemophagocytes were seen in the alveoli and lymph nodes from monkey #5417, which showed severe anemia at 7 days p.i. (S10A Fig). Diffuse eosinophilic and plasma cell infiltration was seen in the mesenteric lymph nodes, small intestines and large intestine from monkey #5412, which showed prolonged viral excretion after initial infection (S10B Fig). No viral antigens were detected in extrapulmonary tissues.

## Discussion

In our previous study of the SARS-CoV HKU39849 isolate, we inoculated six cynomolgus monkeys via the intranasal, intragastric, intravenous, or intratracheal routes and found that only intratracheal inoculation with 10^8^ TCID_50_ virus in 5 mL of medium induced acute pneumonia [31]. A low dose (10^3^ TCID_50_ in 3.5 mL) administered intranasally failed to establish an infection, whereas a high dose (10^6^ TCID_50_ in 3.5 mL) succeeded; indeed, infection was detected in nasal and throat swabs within 7 days post-inoculation. In addition, an epidemiological study suggests that COVID-19-associated conjunctivitis is a possible transmission route for SARS-CoV-2 [32]. Therefore, in this study we used a combined inoculation protocol comprising the nasal, intratracheal, and conjunctive routes, and used a high titer of SARS-CoV-2. Shed virus was detected in the upper respiratory and intestinal tracts of infected monkeys, but not consistently (even in nasal and throat swabs); however, one monkey from the CD3+ low group showed prolonged shedding of virus in rectal swabs. Other macaque models infected with clinical isolates of SARS-CoV-2 show similar results [20, 33]. One of these studies showed that viral RNA levels in throat and nasal swabs from young cynomolgus monkeys peaked at Day 1 or 2 post-inoculation; however, they peaked at Day 4 in older monkeys [20]. A few conjunctival swab samples were positive for viral RNA, but not for subgenomic mRNA; in addition, none of the monkeys developed obvious conjunctivitis during the observation period in this study. Taken together, these data suggest that a combination of the intranasal and intratracheal routes (at least) might be appropriate for vaccine studies. In addition, a previous study suggests that the presence of subgenomic mRNA in throat and/or nasopharyngeal swabs should be considered when testing vaccine efficacy [34]. Our own study using young adult cynomolgus monkeys suggests that peripheral T lymphocytes (CD3+) are associated with pneumonia severity. Thus, it is important to consider both the age of the individual and T cell population when selecting animals for vaccine studies [18].

Peripheral blood lymphocyte subsets in humans are affected by factors such as gender, age, and ethnicity, and by lifestyle factors such as stress [35]. In this study, we used young healthy monkeys, which showed a wide range of peripheral CD3+ cells. The immune system of non-human primates may also be affected by environmental and physiological conditions [36, 37].

According to an epidemiological study of COVID-19, about 10% of the global population may be infected by October 2020; however, most infected people are asymptomatic or mildly symptomatic [38]. That said, some people develop severe pneumonia resulting in respiratory failure, sepsis, and even death (the current fatality rate is 0.15–0.20%). Similar to SARS-CoV and MERS-CoV, older age is a risk factor for severe SARS-CoV-2. Whereas absolute lymphopenia is not specific to COVID-19, low CD3+, CD4+, and CD8+ T cell counts in peripheral blood have been observed in severe cases of COVID-19 [39]. These cases also present with comorbidities such as chronic underlying diseases. Zheng et al. reported that the total CD3+ count is lower in both mild and severe cases of COVID-19 than in healthy controls, but that CD3+, CD8+, and NK cell counts are significantly lower in severe cases [40]. In addition, functional exhaustion (e.g., reduction of CD107a expression and IFN-γ, IL-2, and TNF-α production by CTLs and NK cells) occurred in severe cases.

Murine models of SARS-CoV and MERS-CoV infection suggest that failure to induce an early IFN-I response leads to severe pathology and disease [41, 42]. Sera from hospitalized COVID-19 patients show reduced IFN-I and -III levels in response to SARS-CoV-2, but a significant increase in inflammatory chemokines and cytokines [43]. In the current study, transcriptome analysis revealed that innate anti-viral immune responses occurred during the early phase of infection in both the CD3+ high and low groups. In both groups, IRF2, which regulates type I IFN production, was activated during the early phase of infection and upon re-infection. However, in the CD3+ low group, inflammation overwhelmed the T cell response. This is supported by the kinetics of T cell-associated cytokine and chemokine production in monkey sera. Thus, a strong inflammatory response, coupled with a weak/delayed T cell response, was critical for the development of more severe SARS-CoV-2 in the CD3+ low group. By contrast, an early type I IFN-related innate immune response controlled viral replication and dispersion at an early stage in the CD3+ high group.

On Days 7–10 after the initial inoculation, two monkeys from the CD3+ low group became lethargic, with decreased hemoglobin levels and RBC counts suggestive of severe anemia. In some cases of COVID-19, low hemoglobin levels indicate anemia [2, 44-46]. The mechanism underlying anemia in COVID-19 patients is unclear; however, virus infection and inflammation impact iron metabolism [47-49]. Levels of serum ferritin, an intracellular protein that maintains iron levels, mirror the degree of inflammation in infectious diseases. In this study, we did not measure ferritin levels in blood from infected monkeys; however, studies show that hospitalized COVID-19 patients have high ferritin levels [2, 46]. The impact of anemia and high ferritin levels on outcome after SARS-CoV-2 infection is unclear [45]. In this study, one of two monkeys (#5417) with anemia that was sacrificed for planned autopsy showed severe acute pneumonia and hemophagocytes in the cervical lymph nodes. Another (#5412) showed extreme lethargy and anemia on Day 10; however, the monkey ate a piece of apple despite showing loss of appetite. Therefore, we continued to observe this animal until recovery within 14 days p.i., at which point seroconversion occurred. Monkey #5412 showed a low clinical score and excreted infectious virus from intestine for 3 weeks.

Pathological evaluation revealed varying degrees of virus infection and host response in the lungs of SARS-CoV-2-infected monkeys at 7 days p.i. Morphologically, SARS-CoV-2 replicated in epithelial cells in the pulmonary bronchus and alveoli of monkey #5404, resulting in mild pneumonia. Similar to SARS-CoV infection, expression of ACE2 and SARS-CoV antigen-positive cells did not overlap [50]. In a severe case (monkey #5417), pulmonary edema was observed, suggesting severe damage to pneumocytes. The pathological features were early stage diffuse alveolar damage, with hyaline membranes and a few multinucleated giant cells, similar to human cases of SARS and COVID-19 [7, 9, 11, 51-53]. Activated macrophages rather than lymphocytes were seen in the alveoli of monkey #5417, suggesting that massive inflammatory reactions were induced in the lungs. Lack of an active immune response and epithelial regeneration results in a poor outcome [51]. In this study, we used young monkeys; many regenerated type II cells were seen in the lungs of monkey #5417, and high levels of seroconversion occurred in monkey #5412, even from CD3+ low groups.

Most infected people are asymptomatic or show mild symptoms during SARS-CoV-2 infection; thus some researchers wonder whether SARS-CoV-2 infection triggers protective immunity against re-infection [54]. A rhesus macaque model clarified that SARS-CoV-2 infection results in protective immunity against re-infection [55]. The latest study reporting human cases of COVID-19 indicate that the neutralizing antibodies against SARS-CoV-2 last only for a few months [56]. The results of the present study suggest the magnitude of neutralizing antibody titers in infected monkeys is dependent on disease severity, similar to human cases [56]. In addition, these monkeys developed a rapid immune response against a second infection with another challenge strain. NK cell and IL-17 responses, suggesting involvement of Th17 cells, were stronger after the second infection than after the initial infection. Transcriptome analysis revealed that upregulation of innate immune responses, rather than T and B cell responses, in the CD3+ low group contributed to a marked reduction in viral replication and less severe pathology, even after a second infection. Seroconversion in monkeys is common after acute virus infections; indeed, we found virus-specific IgM, IgG, and IgA antibodies in the sera. IgM antibodies appeared together with IgG and, later, IgA; however, titers decreased within 3 weeks after inoculation. This result is similar to that of a human cohort study reporting co-induction of IgM and IgG during SARS-CoV infection [57]. SARS-CoV-2-specific IgG antibodies are predominantly specific for the S-/RBD- and N proteins. IgG levels in symptomatic groups are significantly higher than those in asymptomatic groups during the acute phase [58].

Asymptomatic cases also show lower levels of pro- and anti-inflammatory cytokines. Similar to human cases of COVID-19, our monkeys showed different immune responses and even seroconversion. After the second inoculation, all monkeys generated high titers of virus-specific IgA and IgG, suggesting re-infection.

In this study, we used two clinical isolates of SARS-CoV-2, one from East Asia and one from Europe. After identification of the first case of COVID-19 in Japan on January 15, 2020, an epidemiological study of the SARS-CoV-2 genome revealed that the primary clusters identified in January and February in Japan were related to the Wuhan-Hu-1 isolates from China [59]. Soon after the primary wave from China, we faced a second wave of COVID-19 cases caused by lineages imported by returnees from Europe and North America. Thus, we based the infection experiments in this study on the current situation in Japan. We found that previous infection with a Wuhan-Hu-1-related isolate of SARS-CoV-2 led to a less severe illness upon re-infection with a heterologous strain (an S-G614 variant from Europe).

We also determined the mutation patterns in SARS-CoV-2 isolates from the lung of monkey #5412 at 6 weeks after the initial inoculation. The most common base changes were C > T, which were synonymous variants in the ORF1ab region of the monkey isolate. This nucleotide substitution is common in SARS-CoV-2 genomes isolated from humans [60, 61]. C > T transitions are thought to be induced by cytosine deaminases [60].

Taken together, the data presented herein suggest that a low CD3+ T cell count in peripheral blood might be an important risk factor for more severe COVID-19. We acknowledge that the study has some limitations; the small number of monkeys (due to ethical reasons) in particular. However, the data suggest that the peripheral T lymphocyte population is associated with severity of pneumonia caused by SARS-CoV-2 infection.

## Materials and methods

### Ethical statements

All animal experiments complied with Japanese legislation (Act on Welfare and Management of Animals, 1973, revised in 2012) and guidelines under the jurisdiction of the Ministry of Education, Culture, Sports, Science and Technology, Japan (Fundamental Guidelines for Proper Conduct of Animal Experiment and Related Activities in Academic Research Institutions, 2006). Animal care, housing, feeding, sampling, observation, and environmental enrichment were performed in accordance with these guidelines. Every possible effort was made to minimize suffering. The protocols were approved by the committee of biosafety and animal handling and by the committee of ethical regulation of the National Institute of Infectious Diseases, Japan (authorization nos. 519004-I, -II, and -III for monkey experiments; authorization no. 119176 for rabbit immunizations). Each monkey was housed in a separate cage at the National Institute of Infectious Diseases, Japan, an all received standard primate feed and fresh fruit daily, and had free access to water. Each rabbit was housed in a separate cage at the National Institute of Infectious Diseases, Japan, and all received standard rabbit feed and had free access to water. Animal welfare was observed on a daily basis. Inoculation of monkeys with virus was conducted under ketamine-xylazine anesthesia (intramuscular injection of a mixture of 50 mg/mL ketamine and 20 mg/mL xylazine [2:1; 0.2 mL/kg]). Sampling procedures were conducted under anesthesia (10 mg/kg ketamine; intramuscular injection). Monkeys were sacrificed under excess anesthesia with ketamine (intramuscular injection). Rabbits were sacrificed under excess anesthesia with pentobarbital sodium (64.8 mg/kg intravenous injection).

### Biological safety

All work with SARS-CoV-2 was conducted under biosafety level-3 (BSL-3) conditions in the National Institute of Infectious Diseases, Japan. All experimental animals were handled in a biosafety level 3 animal facility in accordance with the guidelines of this committee (approval no. 19-60, 20-1). Animals were contained in a glovebox system in the ABSL-3 facility during experimental infection. All personnel used respiratory protection when handling infectious samples (respirator type N95). Surface disinfection was performed using 80% ethanol, while liquids, solid waste, cages, and animal wastes were steam sterilized in an autoclave.

### Cells and viruses

VeroE6/TMPRSS2 cells and SARS-CoV-2 human isolates were kindly prepared and provided by Dr. Shutoku Matsuyama and Dr. Makoto Takeda (Department of Virology III, National Institute of Infectious Diseases, Japan) [62]. Cells were cultured in Dulbecco’s modified Eagle’s medium (DMEM, low glucose (Sigma-Aldrich, St. Louis, MO)) containing 5% fetal bovine serum (FBS), 50 IU/mL penicillin G, and 50 μg/mL streptomycin (5DMEM). The virus strains used in this study are shown in Table 1. Stocks of the 2019-nCoV/Japan/TY/WK-521/2020 isolate (refer as WK-521) of SARS-CoV-2 (accession no. EPI_ISL_408667) and the hCoV-19/Japan/QH-329-037/2020 isolate (refer as QH-329-037) were propagated eight times or twice, respectively, and titrated on VeroE6/TMPRSS2 cells in DMEM containing 2% FBS (2DMEM).

Whole-genome amplification of strain QH-329-037 was carried out using the modified version of ARTIC Network’s protocol for SARS-CoV-2 genome sequencing by replacing some of the primers for multiplex PCR [63]. A next generation sequencing (NGS) library was constructed using the QIAseq FX DNA library kit (Qiagen, Hilden, Germany) and sequenced using the NextSeq 500 platform (Illumina, San Diego, CA). NGS reads were mapped to the SARS-CoV-2 Wuhan-Hu-1 reference genome sequence (GenBank accession no. MN908947) using bwa mem [64], followed by trimming the primer region by “trim_primer_parts.py” (https://github.com/ItokawaK/Alt_nCov2019_primers). For determination of the nearly full-length genome sequence, the trimmed reads were assembled using A5-miseq v.20140604 [65]. The full genome sequence of strain QH-329-037 has been deposited in the Global Initiative on Sharing All Influenza Data database (GISAID) under accession ID EPI_ISL_529135.

To eradicate mycoplasma contamination, cells and strain WK-521 were treated with an anti-mycoplasma reagent, MC-210 (0.5 µg/mL; Waken, Kyoto, Japan). Mycoplasma contamination was confirmed by PCR using the TaKaRa PCR Mycoplasma Detection Set (Takara, Shiga, Japan).

### Animal experiments

Twenty-five female adult cynomolgus macaques (*Macaca fascicularis*) imported from China were purchased from Hamri Co., Ltd (Ibaraki, Japan) in 2018 and maintained in the animal facility of the National Institute of Infectious Diseases, Japan. At around 4 weeks before experimental infection, blood samples were collected from all animals under anesthesia with ketamine (intramuscular injection) (Fig. 1A). Sera were used for neutralization assays against SARS-CoV-2. Ethylenediaminetetraacetic acid (EDTA) blood samples were used for hematologic tests and flow cytometry analysis. Six monkeys (young adult females, 5 years old) were selected for experimental infection with SARS-CoV-2. At 14 days before inoculation with the virus, a small implantable thermo logger (DST micro-T: 8.3 × 25.4 mm; Star-oddi, Gardabaer, Iceland) was set intraperitoneally under ketamine anesthesia. The loggers were retrieved at necropsy. Six monkeys were transferred to the animal facility at biosafety level 3 and allowed to acclimatize for 1 week. The animals were observed daily for clinical signs (dietary intake, including pellets and fruits, drinking, attitude in front of regular observers, and stool consistency) using a standardized scoring system until the end of the study. Scoring was performed as follows: daily intake of pellets (0–5), fruits including orange and apple (0–5), and drinking water (0–5), attitude in front of regular observers (i.e., standing up, show interest in the outside, getting attention, intimidation, up and down movement: 0–5), stool consistency (color, stiffness, form, volume, frequency: 0–5). The total score was the sum of all five component scores.

The six monkeys were anaesthetized by intramuscular injection of a mixture of 50 mg/mL ketamine and 20 mg/mL xylazine (2:1; 0.2 mL/kg). After collecting samples, including blood and swabs, monkeys were inoculated with an isolate of SARS-CoV-2 (WK-521) via the intranasal (0.125 mL, sprayed into the right nostril; Keytron, Ichikawa, Japan), conjunctival (0.1 mL dropped into the right eye), and intratracheal (1 mL of virus solution plus 2 mL of saline via a catheter; 6Fr; Atom Medical, Tokyo, Japan) routes (all three routes combined). On Days 0, 1, 4, 7, 10, 14, 21, 28, and 35 after initial virus inoculation, clinical samples (conjunctiva, nasal, throat, and rectal swabs, and blood samples) were collected after monkeys were weighed under anesthesia. Two animals were euthanized at 7 days post-initial inoculation, and four animals were re-inoculated with another isolate of SARS-CoV-2 (QH-329-037) at 35 days p.i. After re-inoculation, two animals were euthanized at 7 days post-second inoculation (R7 days p.i.), and the remaining two were euthanized at R14 days p.i. Clinical samples were collected at R1, R4, R7, R10, and R14 days p.i.

### Virus titration

Tissue samples in Lysing Matrix tubes containing beads (Lysing Matrix A; MP Biomedicals, Irvine, CA) were homogenized using a mini Bead-Beater (Biospec Products, Bartlesville, OK) at 100 rpm for 30 sec (twice), and then diluted in 2×DMEM to yield 10% homogenates. After centrifugation at 10,000 × *g* for 1 min at 4℃, the supernatants were used for titration on VeroE6/TMPRSS2 cells. Swab samples were also used for titration. Inoculated cells were assessed for CPE at 5 days p.i. The detection limit was 10^1.5^ TCID_50_/mL 10% tissue homogenate or swab sample.

### Real-time RT-PCR of SARS-CoV genome and detection of viral sequence

Total RNA was extracted from 100 µL swab samples, tissue homogenates, or blood samples using a TRIzol™ Plus RNA Purification Kit (Thermo Fisher Scientific, Waltham, MA) and used to quantify the SARS-CoV-2 genome. On-column PureLink DNase (Thermo Fisher Scientific) treatment was performed during RNA purification, and RNA samples were dissolved in 30 µL RNase-free water. The viral RNA copy number in samples from monkeys was estimated by real-time RT-PCR [66]. Subgenomic viral RNA transcripts were also detected in N gene transcripts. The primer and probe sets are shown in S4 Table. Real-time RT-PCR was performed using the QuantiTect Probe RT-PCR Kit (QuantiTect, Qiagen, Venlo, Netherlands) and a LightCycler 480 (Roche, Basal, Switzerland) or Mx3005P (Stratagene, La Jolla, CA) apparatus. The thermal cycling conditions were as follows: 50°C for 30 min, 95°C for 15 min, and 45 cycles at 95°C for 15 s and 60°C for 1 min (N2 primer and probe set); or 50°C for 30 min, 95°C for 15 min, and 40 cycles of 94°C for 15 s and 60°C for 1 min (N1 set and the sgRNA transcript primer and probe sets).

Some samples containing high viral RNA copy numbers were sent for viral sequence analysis by gene analysis services (Takara Bio, Shiga, Japan). The next generation sequencing (NGS) library was prepared using the SuperScript IV First-Strand Synthesis System (Thermo Fisher Scientific), Q5 Hot Start DNA Polymerase (New England Biolabs, Ipswich, MA), and the QIAseq FX DNA Library Kit (Qiagen). The viral genome region was amplified specifically by multiplex PCR [63], and the entire sequence of the viral genome was obtained using the next generation sequencer MiSeq (Illumina, San Diego, CA) with a read length of 250 nt. FASTQ data were imported into the CLC Genomics Workbench (version 11, Qiagen), and the sequence reads were aligned to the reference sequence Wuhan-Hu-1 (accession no. MN908947.3). The threshold variant frequency was 50%. The amino acid substitutions were analyzed on NextClade (https://clades.nextstrain.org/). Genome sequences were deposited in the DNA Data Bank of Japan (DDBJ) (https://www.ddbj.nig.ac.jp/index.html).

### Hematological analysis

Complete blood cell counts, hematocrit, and hemoglobin levels in peripheral blood collected in EDTA tubes were measured by an autoanalyzer (VetScan HM2; ABAXIS, Union City, CA). Neutrophil, lymphocyte, monocyte, eosinophil, and basophil counts were measured by microscopic analysis. Blood biochemistry (Glob, ALB, glucose, alkaline phosphatase (ALP), and blood urea nitrogen (BUN)) of lithium-heparin treated whole blood samples was analyzed using the VetScan VS2 (ABAXIS).

### Flow cytometric analyses

Flow cytometry analysis was conducted to determine the number of T, B, NK, CD4+, and CD8+ cells in peripheral blood samples from monkeys. Cell staining was performed using the NHP T/B/NK Cell Cocktail (Becton Dickinson (BD) Company, Franklin Lakes, NJ) and the NHP T Lymphocyte Cocktail (BD), according to the manufacturer’s instructions. After treatment with BD FACS lysing solution (BD), samples were analyzed by flow cytometry using a BD FACSCanto II analyzer (BD). Flow cytometry data were analyzed using FlowJo software (v10.7.1, FlowJo LLC, Ashland, OR).

### Histopathology and immunohistochemistry

Animals were euthanized by exsanguination under excess ketamine anesthesia and then necropsied. Tissue samples were immersed in 10% phosphate-buffered formalin, embedded in paraffin, sectioned, and stained with hematoxylin and eosin. Immunohistochemical analysis was performed using a polymer-based detection system (Nichirei-Histofine Simple Stain Human MAX PO®; Nichirei Biosciences, Inc., Tokyo, Japan). Antigen retrieval from formalin-fixed monkey tissue sections was performed by autoclaving in retrieval solution (pH 6.0; Nichirei Biosciences) at 121°C for 10 min. Hyper-immune rabbit serum raised against the GST-tagged N protein of SARS-CoV-2 (produced in-house) was used as the primary antibody to detect viral antigens. Peroxidase activity was detected with 3,3’-diaminobenzidine (Sigma-Aldrich, St. Louis, MO). Hematoxylin was used for counterstaining. The polyclonal antibody against GST-tagged N protein of SARS-CoV-2 was prepared as follows: first, the recombinant N protein was constructed by inserting the N gene of SARS-CoV-2 into the pGEX-6P vector (GenScript Japan, Tokyo, Japan). Next, the amino acid sequence was optimized to the bacterial codon. The vector was then used to transform *Escherichia coli* strain BL21 (Takara Bio, Shiga, Japan). Expression of the GST-N protein of SARS-CoV-2 was induced by isopropyl-D-1-thiogalactopyranoside (0.3 mM IPTG, Takara Bio). The cell pellets were sonicated, and the inclusion bodies containing the fusion protein were collected. The fusion proteins were extracted from SDS-PAGE gels after reverse staining (AE-1310 EzStain Reverse, Atto, Tokyo, Japan), concentrated using a spin column (Pall centrifugal device 0.2 µm, Pall Corporation, Port Washington, NY), and diluted in PBS using Amicon Ultra-0.5mL Centrifugal Filters (Ultracel-50k). Two New Zealand White rabbits (1.5 kg < body weight; female; SLC, Shizuoka, Japan) were immunized (four times at 2-week intervals) with the purified protein conjugated to TiterMax Gold (Sigma-Aldrich). Rabbits were sacrificed under excess anesthesia with pentobarbital sodium (64.8 mg/kg), and whole blood was collected by cardiac puncture using an 18 G needle. After separating sera by centrifugation, IgG was purified from the rabbit serum using a Melon Gel IgG Spin Purification Kit (Thermo Fisher Scientific) and then used for immunohistochemistry.

### Neutralization assay

During the observation period, blood was obtained under anesthesia with ketamine. Serum samples were collected by centrifugation and inactivated by heating to 56°C for 30 min. Serum samples were titrated (in duplicate) from 1:10 to 1:1280 in 96-well plates and reacted with 100 TCID_50_ of SARS-CoV-2 (WK-521 or QH-329-037) at 37°C for 1 h before addition of VeroE6/TMPRSS2 cells. Cells were incubated at 37°C for 5 days and examined twice for evidence of viral CPEs. The neutralizing antibody titer was determined as the reciprocal of the highest dilution at which no CPE was observed.

### ELISAs

To assess the specificity of the IgM, IgA, and IgG antibodies produced by the infected monkeys, recombinant SARS-CoV-2 trimeric spike, RBD, or nucleocapsid protein were used as antigens in ELISAs. Briefly, 96-well assay plates (Corning Inc., Corning, NY) were coated overnight at 4℃ with 50 ng recombinant protein in coating buffer (pH 9.6). The serum samples were serially diluted (4-fold from 1:400 to 1:409600) in 5% skim milk in PBS (pH 7.2) containing 0.05% Tween 20 (Sigma-Aldrich) (PBS-T). The well contents were discarded and diluted serum samples were added to the plate. After incubation for 1 h at 37℃, the plate was washed three times with PBS-T. The wells were then incubated with an HRP-conjugated goat anti-monkey IgM antibody (KPL #5220-0334, SeraCare Life Sciences, Inc. Milford, MA, 1/5000, 50 µL/well), an HRP-conjugated goat anti-monkey IgA antibody ((KPL #5220-0332, SeraCare Life Sciences, 1/5000, 50 µL/well), or an HRP-conjugated goat anti-monkey IgG heavy and light chain antibody (A140-102P, 1/10000, 50 µL/well, Thermo Fisher Scientific) in 5% skim milk in PBS-T for 1 h at 37℃. After three washes with PBS-T, an ABTS substrate (Roche, Basel, Switzerland) was added to the wells, and the plates were incubated for 30 min at room temperature. The optical density (OD) of each well was measured at 405 nm using a microplate reader (Model 680, Bio-Rad). The mean OD value plus three standard deviations (2 × mean + 3 × SD) was calculated using serum samples from pre-infected monkeys and was used as the cut-off for the Ig ELISAs.

### Detection of inflammatory cytokines and chemokines

All serum samples tested in the BSL2 laboratory (all of which were confirmed negative for viral RNA by RT-PCR) were irradiated for 1 min with UV-C light. Cytokine and chemokine levels in monkey sera were measured using a MILLIPLEX MAP Non-Human Primate Cytokine Magnetic Bead Panel - Premixed 23 Plex - Immunology (Milliplex MAP kit, Merck Millipore, Burlington, MA), which includes the following 23 cytokines and chemokines: G-CSF, granulocyte macrophage colony-stimulating factor (GM-CSF), interferon gamma (IFN-γ), IL-1ra, IL-1β, IL-2, IL-4, IL-5, IL-6, IL-8, IL-10, IL-12/23 (p40), IL-13, IL-15, IL-17, IL-18, monocyte chemotactic protein-1 (MCP-1), macrophage inflammatory protein 1 alpha (MIP-1α), MIP-1β, sCD40L, transforming growth factor alpha (TGF-α), tumor necrosis factor alpha (TNF-α), and vascular endothelial growth factor. The assay samples were read on a Luminex 200™ instrument with xPONENT software (Merck Millipore), as described by the manufacturer.

### RNA sequencing and data analyses

Whole blood was collected from animals at multiple time points using PAXgene Blood RNA Tubes (PreAnalytiX, Hombrechtikon, Switzerland). Tubes were frozen at −80°C until RNA extraction. RNA was extracted using PAXgene Blood RNA Kits (PreAnalytiX) and shipped to Macrogen Corp. Japan (Kyoto, Japan) for NGS sequencing. Next, cDNA libraries were prepared using a TruSeq Stranded Total RNA LT Sample Prep Kit (Illumina) in accordance with the TruSeq Stranded Total RNA Sample Prep Guide (Part #15031048 Rev. E protocol). Next, the cDNA libraries were paired-end sequenced (read length = 101 bp) on a NovaSeq6000 sequencer (Illumina). Raw FASTQ files were quality checked using fastqc v0.11.8 [67], and low-quality bases from paired reads were trimmed using Trimmomatic v0.39 [68]. Paired reads were aligned to the Macaca fascicularis genome (version 5.0, Ensembl release 101) using the STAR aligner v2.7.3a [69] and default settings. Read fragments (paired reads only) were quantified per gene per sample using featureCounts v1.6.0 [70]. All raw RNA seq fastq files were uploaded to the DDBJ Sequence Read Archive (DRA accession number: DRA010881). All functional analyses of transcriptomic data were performed in the R statistical environment (v3.6.2). Significantly differentially expressed genes between samples collected before and after virus infection were identified using DESeq2 v1.26.0 [71] with default settings, and a minimum adjusted *P*-value significance threshold of 0.05. Volcano plots were created from shrunken log2-fold change values for each gene, calculated by DESeq2 (shrinkage type: normal). For the heatmaps, DESeq2-normalized counts per gene were plotted using the heatmap package [72]. Gene set enrichment analyses (GSEA) were conducted using tmod v0.44 [73], with count data normalized with the voom function within the limma package v3.42.2 [74]. GSEA (with default settings and a minimum *P*-value significance threshold of 0.01) was conducted between samples collected from animals in the CD3+ high and low groups after virus infection, and samples collected before and at multiple time points after virus infection.

### Statistical analysis

Data are expressed as the mean and standard error of the mean. Statistical analyses were performed using Graph Pad Prism 8 software (GraphPad Software Inc., La Jolla, CA). Intergroup comparisons (i.e., changes in clinical scores, blood analysis results, and cytokine levels) were performed using Sidak’s multiple comparisons test after application of mixed-effects models for repeated measures analysis. The correlation coefficient was evaluated by Spearman’s correlation analysis of the neutralization and ELISA test results. A *P*-value of <0.05 was considered statistically significant.

## Acknowledgements

We thank Dr Shutoku Matsuyama and Dr Makoto Takeda (National Institute of Infectious Disease) for providing VeroE6-TMPRSS2 cells and SARS-CoV-2 isolates. We also thank Dr Masayuki Shimojima, Dr. Hideki Asanuma, Dr Makoto Kuroda, Dr Takushi Nomura, Dr Hiroyuki Yamamoto, Dr Tetsuro Matano, Dr Shinji Watanabe (National Institute of Infectious Diseases), Dr Shintaro Shichinohe, and Dr Kensuke Nakajima (Nagasaki University, Nagasaki, Japan) for helpful discussion. We also thank our colleagues at the Institute, especially Ms Midori Ozaki, Ms Takiko Yoshida, Dr Michiyo Kataoka, Dr. Dai Izawa, and Ms Yuriko Suzaki, for technical assistance.

## Data availability

All relevant data are provided in the manuscript and the Supporting Information files.

## Author contributions

Conceptualization: NN, NI-Y, T. Su, HH

Data Curation: NN

Formal Analysis: NN, KS, AA

Funding Acquisition, NN, HS, T. Su, HH

Investigation: NN, NI-Y, KS, AA, NS, MS, NK, TA, YS, TH, YK, YA, SI, HK, SF, T. Se, T. Su

Methodology: NN, NI-Y, KS, AA, YA, T. Su

Project Administration: NN, T. Su, HH

Resources: NN, NI-Y, KS, AA, HS, T. Su, HH

Supervision: NN, T. Su, HH

Validation: NN, T. Su, HH

Visualization: NN, KS

Writing - Original Draft: NN, KS

Writing - Review & editing: NN, NI-Y, KS, AA, NS, MS, NK, TA, YS, TH, YK, YA, SI, HK, SF, T. Se, HS, T. Su, and HH

## Supporting information

**S1 Fig. Selection of monkeys for experimental infection.** (A) Body weight of the 25 monkeys in Figure 1. (B) Analysis of lymphocytes in peripheral blood from 15 animals weighing <3.4 kg. Each dot represents data from an individual animal. The blue and red colored symbols denote data from the CD3+ high and low groups, respectively.

**S2 Fig. Clinical course in cynomolgus monkeys inoculated with SARS-CoV-2.** Body weight was measured under anesthesia at various time points after inoculation (A). Biochemical markers including globulin (Glob), albumin (ALB), glucose, alkaline phosphatase (ALP), and blood urea nitrogen (BUN) in lithium-heparin treated whole blood samples were measured at various time points after inoculation (B). Six cynomolgus monkeys were used. Cool (blue and aqua) and warm (red and orange) colored symbols and lines indicate data from the CD3+ high and CD3+ low groups, respectively. Each dot/line represents data from an individual animal. The brown dashed line on the vertical axis indicates the day of second inoculation.

**S3 Fig. Variations in deep body temperature detected by the temperature logger.** Thermo logger probes were set intraperitoneally at 14 days before inoculation. Black arrows, animal transfer date (under anesthesia) from the animal facility to the animal biosafety level 3 (ABSL3); red and yellow arrow heads, virus inoculation under anesthesia with a mixture of ketamine and xylazine; Red brace, deviation from diurnal variation indicates high fever. The fluctuation of deep body temperature within a day was maintained during ABSL3 acclimatization. A drop in deep body temperature due to the mixed anesthesia was observed on the day of inoculation.

**S4 Fig. Hematological examination of cynomolgus monkeys inoculated with SARS-CoV-2.** Erythrocyte analysis, including total red blood cells (RBC) and hematocrit (HCT), was performed using EDTA-treated whole blood samples taken at various time points after inoculation (A). Absolute white blood cell (WBC) count, including total WBC, neutrophils, eosinophils, and basophils, in EDTA-treated whole blood samples was measured at various time points after inoculation (B). Markers of leukocyte differentiation, CD4 and CD8, were detected by flow cytometry at various time points after inoculation (C). Cool (blue and aqua) and warm (red and orange) colored symbols and lines indicate data from the CD3+ high and CD3+ low groups, respectively. Each dot/line represents data from an individual animal. The brown dashed line on the vertical axis indicates the day of the second inoculation.

**S5 Fig. Cytokine and chemokine levels in serum samples from cynomolgus monkeys inoculated with SARS-CoV-2.** Sera were obtained from six monkeys at various time points after inoculation. Pro-inflammatory cytokines and chemokines (A), helper T cell-related cytokines (B), and other representative factors in serum that drive proliferation of epithelial cells (TGF-α) and neutrophils (IL-8) (C) were profiled by multiplex analysis. Assays were performed using unicate samples per time point. Cool (blue and aqua) and warm (red and orange) colored symbols and lines indicate data from the CD3+ high and CD3+ low groups, respectively. Each dot/line represents data from an individual animal. The brown dashed line on the vertical axis indicates the day of the second inoculation.

**S6 Fig. Detection of subgenomic RNA in clinical samples and tissue samples from cynomolgus monkeys inoculated with SARS-CoV-2.** Virus RNA-positive samples from Figures 3 and 6 were re-examined to detect viral RNA and subgenomic RNA using three primer sets (A and B, respectively).

**S7 Fig. Transcriptome analysis of blood samples from cynomolgus monkeys inoculated with SARS-CoV-2.** Volcano plot showing the magnitude and significance of differentially expressed genes between samples collected from animals before (Day 0) and after (Days 1, 4, 7, R0, R1, R4, and R7) virus infection (A). Red plots indicate genes that were upregulated significantly (331 genes) after virus infection, and blue plots indicate genes that were downregulated significantly (176 genes) after virus infection (adjusted *P*-value < 0.05). Plots shown in brighter red or blue represent genes that were either upregulated (190/331 genes) or downregulated (86/176 genes) by more than 2-fold. Expression of immunity-related genes in peripheral whole blood samples collected from animals before (Day 0) and after (Days 1, 4, 7, R0, R1, R4, and R7) virus infection (B). Heatmaps showing normalized counts per gene, scaled by rows of 78 immune-related genes among the 507 genes significantly upregulated or downregulated by virus infection (adjusted *P*-value < 0.05). Gene symbols are listed on the right. Yellow and green/red dots indicate genes related to innate immunity and inflammation, respectively. Each column represents a different sample. Animal ID, days post-virus infection (dpi), and CD3+ expression in each sample are shown at the top.

**S8 Fig. Double immunohistochemistry to detect virus antigens (brown) and ACE2 (green) in the lungs at 7 days after initial inoculation.** ACE2 was detected in the intact brush border of the respiratory epithelia in the intrapulmonary bronchus (black arrows); however, no cells were positive for viral antigens (red arrow; upper row, left). Viral antigen was detected in linear pneumocytes and type I pneumocytes (red arrow), and slight expression of ACE2 was detected in round pneumocytes (suggestive of type II pneumocytes) (black arrows), in the alveolar area in the absence of inflammatory infiltration (upper row, right). Strong expression of ACE2 on large pneumocytes suggested hyperplasia of type II pneumocytes (black arrows, lower row); there were no degenerated viral antigen-positive pneumocytes (red arrow, lower row, left) at the lesion sites in the alveolar area. Bars, 20 µm. An anti-SARS-CoV-2 nucleocapsid protein rabbit polyclonal antibody and an anti-ACE2 goat-polyclonal antibody were used for IHC.

**S9 Fig. Lung pathology in cynomolgus monkeys receiving a second inoculation with SARS-CoV-2.** Representative histopathology images of lungs from monkeys obtained at 7 days (#5403 and #5412) or 14 days (#5399 and #5405) after re-infection with QH-329-037 strain. Cellular infiltration, including lymphocytes and macrophages, can be seen around the bronchi and in the alveoli in the middle lobe of the right lung from monkey #5403 (first row). Lymphoid aggregates, including alveolar macrophages, were observed in the alveoli in the upper lobe of the right lung from monkey #5399 (second row). Lymphoid aggregates around small vessels (red arrowheads) and fibrotic inflammation with lymphocyte aggregation in the alveolar area and pleura (blue arrowheads) were seen in the right lung from monkeys #5412 and #5405 (third and fourth rows). Scale bars: 500 µm (left column), 50 µm (middle column), and 20 µm (right column). Hematoxylin and eosin staining (H&E).

**S10 Fig. Representative images of histopathological lesions from cynomolgus monkeys after experimental inoculation with SARS-CoV-2.** Representative hemophagocytosis images of the lung and lymph nodes from monkey #5417 obtained at 7 days after infection with WK-521 strain (A). Hemophagocytes are seen in the alveoli and sinus of the cervical and splenic lymph nodes (yellow arrows). Scale bars: 50 µm (left column) and 20 µm (right column). Hematoxylin and eosin (H&E) staining. Eosinophil (yellow arrows) and plasma cell (blue arrows) infiltration into the mesenteric lymph node and intestines from monkey #5412 at 7 days after the second inoculation with QH-329-037 (B). Cellular infiltration, including eosinophils and plasma cells, can be seen in the sinus of the mesenteric lymph node and the lamina propria of the small and large intestine. Scale bars: 500 µm (left column) and 20 µm (right column). H&E staining.

S1 Table. Cross neutralization of two strains of SARS-CoV-2 in monkey sera after experimental infection.

S2 Table. Summary of the results of next generation sequencing of SARS-CoV-2 from tissue samples of experimentally infected monkeys

S3 Table. D614G variants in tissue samples from experimentally infected monkeys

S4 Table. Primer and probe sets used in this study.

## References

1. Zhu N, Zhang D, Wang W, Li X, Yang B, Song J, et al. A Novel Coronavirus from Patients with Pneumonia in China, 2019. N Engl J Med. 2020;382(8):727–33. Epub 2020/01/25. doi: 10.1056/NEJMoa2001017. PubMed PMID: 31978945; PubMed Central PMCID: PMCPMC7092803.

2. Chen N, Zhou M, Dong X, Qu J, Gong F, Han Y, et al. Epidemiological and clinical characteristics of 99 cases of 2019 novel coronavirus pneumonia in Wuhan, China: a descriptive study. Lancet. 2020;395(10223):507–13. Epub 2020/02/03. doi: 10.1016/S0140-6736(20)30211-7. PubMed PMID: 32007143.

3. Huang C, Wang Y, Li X, Ren L, Zhao J, Hu Y, et al. Clinical features of patients infected with 2019 novel coronavirus in Wuhan, China. Lancet. 2020;395(10223):497–506. Epub 2020/01/28. doi: 10.1016/S0140-6736(20)30183-5. PubMed PMID: 31986264.

4. JHU. COVID-19 Dashboard by the Center for Systems Science and Engineering (CSSE) at Johns Hopkins University (JHU) 2020.

5. Chan JF, Yuan S, Kok KH, To KK, Chu H, Yang J, et al. A familial cluster of pneumonia associated with the 2019 novel coronavirus indicating person-to-person transmission: a study of a family cluster. Lancet. 2020;395(10223):514–23. Epub 2020/01/28. doi: 10.1016/S0140-6736(20)30154-9. PubMed PMID: 31986261.

6. Mizumoto K, Kagaya K, Zarebski A, Chowell G. Estimating the asymptomatic proportion of coronavirus disease 2019 (COVID-19) cases on board the Diamond Princess cruise ship, Yokohama, Japan, 2020. Euro Surveill. 2020;25(10). Epub 2020/03/19. doi: 10.2807/1560-7917.ES.2020.25.10.2000180. PubMed PMID: 32183930; PubMed Central PMCID: PMCPMC7078829.

7. Adachi T, Chong JM, Nakajima N, Sano M, Yamazaki J, Miyamoto I, et al. Clinicopathologic and Immunohistochemical Findings from Autopsy of Patient with COVID-19, Japan. Emerg Infect Dis. 2020;26(9). Epub 2020/05/16. doi: 10.3201/eid2609.201353. PubMed PMID: 32412897.

8. Martines RB, Ritter JM, Matkovic E, Gary J, Bollweg BC, Bullock H, et al. Pathology and Pathogenesis of SARS-CoV-2 Associated with Fatal Coronavirus Disease, United States. Emerg Infect Dis. 2020;26(9). Epub 2020/05/22. doi: 10.3201/eid2609.202095. PubMed PMID: 32437316.

9. Xu Z, Shi L, Wang Y, Zhang J, Huang L, Zhang C, et al. Pathological findings of COVID-19 associated with acute respiratory distress syndrome. Lancet Respir Med. 2020;8(4):420–2. Epub 2020/02/23. doi: 10.1016/S2213-2600(20)30076-X. PubMed PMID: 32085846; PubMed Central PMCID: PMCPMC7164771.

10. Qin C, Zhou L, Hu Z, Zhang S, Yang S, Tao Y, et al. Dysregulation of Immune Response in Patients With Coronavirus 2019 (COVID-19) in Wuhan, China. Clin Infect Dis. 2020;71(15):762–8. Epub 2020/03/13. doi: 10.1093/cid/ciaa248. PubMed PMID: 32161940; PubMed Central PMCID: PMCPMC7108125.

11. Nicholls JM, Poon LL, Lee KC, Ng WF, Lai ST, Leung CY, et al. Lung pathology of fatal severe acute respiratory syndrome. Lancet. 2003;361(9371):1773–8. Epub 2003/06/05. doi: 10.1016/s0140-6736(03)13413-7. PubMed PMID: 12781536; PubMed Central PMCID: PMCPMC7112492.

12. Peiris JS, Chu CM, Cheng VC, Chan KS, Hung IF, Poon LL, et al. Clinical progression and viral load in a community outbreak of coronavirus-associated SARS pneumonia: a prospective study. Lancet. 2003;361(9371):1767–72. Epub 2003/06/05. doi: 10.1016/s0140-6736(03)13412-5. PubMed PMID: 12781535; PubMed Central PMCID: PMCPMC7112410.

13. Wong CK, Lam CW, Wu AK, Ip WK, Lee NL, Chan IH, et al. Plasma inflammatory cytokines and chemokines in severe acute respiratory syndrome. Clin Exp Immunol. 2004;136(1):95–103. Epub 2004/03/20. doi: 10.1111/j.1365-2249.2004.02415.x. PubMed PMID: 15030519; PubMed Central PMCID: PMCPMC1808997.

14. Zhang Y, Li J, Zhan Y, Wu L, Yu X, Zhang W, et al. Analysis of serum cytokines in patients with severe acute respiratory syndrome. Infect Immun. 2004;72(8):4410–5. Epub 2004/07/24. doi: 10.1128/IAI.72.8.4410-4415.2004. PubMed PMID: 15271897; PubMed Central PMCID: PMCPMC470699.

15. Channappanavar R, Perlman S. Pathogenic human coronavirus infections: causes and consequences of cytokine storm and immunopathology. Semin Immunopathol. 2017;39(5):529–39. Epub 2017/05/04. doi: 10.1007/s00281-017-0629-x. PubMed PMID: 28466096; PubMed Central PMCID: PMCPMC7079893.

16. Kim KD, Zhao J, Auh S, Yang X, Du P, Tang H, et al. Adaptive immune cells temper initial innate responses. Nat Med. 2007;13(10):1248–52. Epub 2007/09/25. doi: 10.1038/nm1633. PubMed PMID: 17891146; PubMed Central PMCID: PMCPMC2435248.

17. Lakdawala SS, Menachery VD. The search for a COVID-19 animal model. Science. 2020;368(6494):942–3. Epub 2020/05/30. doi: 10.1126/science.abc6141. PubMed PMID: 32467379.

18. Muñoz-Fontela C, Dowling WE, Funnell SGP, Gsell PS, Balta XR, Albrecht RA, et al. Animal models for COVID-19. Nature. 2020. Epub 2020/09/24. doi: 10.1038/s41586-020-2787-6. PubMed PMID: 32967005.

19. Shi J, Wen Z, Zhong G, Yang H, Wang C, Huang B, et al. Susceptibility of ferrets, cats, dogs, and other domesticated animals to SARS-coronavirus 2. Science. 2020;368(6494):1016–20. Epub 2020/04/10. doi: 10.1126/science.abb7015. PubMed PMID: 32269068; PubMed Central PMCID: PMCPMC7164390.

20. Rockx B, Kuiken T, Herfst S, Bestebroer T, Lamers MM, Oude Munnink BB, et al. Comparative pathogenesis of COVID-19, MERS, and SARS in a nonhuman primate model. Science. 2020;368(6494):1012–5. Epub 2020/04/19. doi: 10.1126/science.abb7314. PubMed PMID: 32303590; PubMed Central PMCID: PMCPMC7164679.

21. Jiang RD, Liu MQ, Chen Y, Shan C, Zhou YW, Shen XR, et al. Pathogenesis of SARS-CoV-2 in Transgenic Mice Expressing Human Angiotensin-Converting Enzyme 2. Cell. 2020;182(1):50–8 e8. Epub 2020/06/10. doi: 10.1016/j.cell.2020.05.027. PubMed PMID: 32516571; PubMed Central PMCID: PMCPMC7241398.

22. Gao Q, Bao L, Mao H, Wang L, Xu K, Yang M, et al. Development of an inactivated vaccine candidate for SARS-CoV-2. Science. 2020;369(6499):77–81. Epub 2020/05/08. doi: 10.1126/science.abc1932. PubMed PMID: 32376603; PubMed Central PMCID: PMCPMC7202686.

23. Chan JF, Zhang AJ, Yuan S, Poon VK, Chan CC, Lee AC, et al. Simulation of the clinical and pathological manifestations of Coronavirus Disease 2019 (COVID-19) in golden Syrian hamster model: implications for disease pathogenesis and transmissibility. Clin Infect Dis. 2020. Epub 2020/03/28. doi: 10.1093/cid/ciaa325. PubMed PMID: 32215622; PubMed Central PMCID: PMCPMC7184405.

24. Kim YI, Kim SG, Kim SM, Kim EH, Park SJ, Yu KM, et al. Infection and Rapid Transmission of SARS-CoV-2 in Ferrets. Cell Host Microbe. 2020;27(5):704–9 e2. Epub 2020/04/08. doi: 10.1016/j.chom.2020.03.023. PubMed PMID: 32259477; PubMed Central PMCID: PMCPMC7144857.

25. Imai M, Iwatsuki-Horimoto K, Hatta M, Loeber S, Halfmann PJ, Nakajima N, et al. Syrian hamsters as a small animal model for SARS-CoV-2 infection and countermeasure development. Proc Natl Acad Sci U S A. 2020;117(28):16587–95. Epub 2020/06/24. doi: 10.1073/pnas.2009799117. PubMed PMID: 32571934; PubMed Central PMCID: PMCPMC7368255.

26. Hartman AL, Nambulli S, McMillen CM, White AG, Tilston-Lunel NL, Albe JR, et al. SARS-CoV-2 infection of African green monkeys results in mild respiratory disease discernible by PET/CT imaging and shedding of infectious virus from both respiratory and gastrointestinal tracts. PLOS Pathogens. 2020;16(9):e1008903. doi: 10.1371/journal.ppat.1008903.

27. Roberts A, Paddock C, Vogel L, Butler E, Zaki S, Subbarao K. Aged BALB/c mice as a model for increased severity of severe acute respiratory syndrome in elderly humans. J Virol. 2005;79(9):5833–8. Epub 2005/04/14. doi: 10.1128/JVI.79.9.5833-5838.2005. PubMed PMID: 15827197; PubMed Central PMCID: PMCPMC1082763.

28. Smits SL, de Lang A, van den Brand JM, Leijten LM, van IWF, Eijkemans MJ, et al. Exacerbated innate host response to SARS-CoV in aged non-human primates. PLoS Pathog. 2010;6(2):e1000756. Epub 2010/02/09. doi: 10.1371/journal.ppat.1000756. PubMed PMID: 20140198; PubMed Central PMCID: PMCPMC2816697.

29. Nagata N, Saijo M, Kataoka M, Ami Y, Suzaki Y, Sato Y, et al. Pathogenesis of fulminant monkeypox with bacterial sepsis after experimental infection with West African monkeypox virus in a cynomolgus monkey. Int J Clin Exp Pathol. 2014;7(7):4359–70. Epub 2014/08/15. PubMed PMID: 25120821; PubMed Central PMCID: PMCPMC4129056.

30. Wolfel R, Corman VM, Guggemos W, Seilmaier M, Zange S, Muller MA, et al. Virological assessment of hospitalized patients with COVID-2019. Nature. 2020;581(7809):465–9. Epub 2020/04/03. doi: 10.1038/s41586-020-2196-x. PubMed PMID: 32235945.

31. Nagata N, Iwata N, Hasegawa H, Sato Y, Morikawa S, Saijo M, et al. Pathology and virus dispersion in cynomolgus monkeys experimentally infected with severe acute respiratory syndrome coronavirus via different inoculation routes. Int J Exp Pathol. 2007;88(6):403–14. Epub 2007/11/28. doi: 10.1111/j.1365-2613.2007.00567.x. PubMed PMID: 18039277; PubMed Central PMCID: PMCPMC2517337.

32. Xia J, Tong J, Liu M, Shen Y, Guo D. Evaluation of coronavirus in tears and conjunctival secretions of patients with SARS-CoV-2 infection. Journal of Medical Virology. 2020;92(6):589–94. doi: 10.1002/jmv.25725.

33. Munster VJ, Feldmann F, Williamson BN, van Doremalen N, Perez-Perez L, Schulz J, et al. Respiratory disease in rhesus macaques inoculated with SARS-CoV-2. Nature. 2020. Epub 2020/05/13. doi: 10.1038/s41586-020-2324-7. PubMed PMID: 32396922.

34. Yu J, Tostanoski LH, Peter L, Mercado NB, McMahan K, Mahrokhian SH, et al. DNA vaccine protection against SARS-CoV-2 in rhesus macaques. Science. 2020;369(6505):806–11. Epub 2020/05/22. doi: 10.1126/science.abc6284. PubMed PMID: 32434945; PubMed Central PMCID: PMCPMC7243363.

35. Choi J, Lee SJ, Lee YA, Maeng HG, Lee JK, Kang YW. Reference values for peripheral blood lymphocyte subsets in a healthy korean population. Immune Netw. 2014;14(6):289–95. Epub 2014/12/22. doi: 10.4110/in.2014.14.6.289. PubMed PMID: 25550695.

36. Nehete PN, Shelton KA, Nehete BP, Chitta S, Williams LE, Schapiro SJ, et al. Effects of transportation, relocation, and acclimation on phenotypes and functional characteristics of peripheral blood lymphocytes in rhesus monkeys (Macaca mulatta). PLOS ONE. 2017;12(12):e0188694. doi: 10.1371/journal.pone.0188694.

37. Rosso MC, Badino P, Ferrero G, Costa R, Cordero F, Steidler S. Biologic Data of Cynomolgus Monkeys Maintained under Laboratory Conditions. PloS one. 2016;11(6):e0157003-e. doi: 10.1371/journal.pone.0157003. PubMed PMID: 27280447.

38. Ioannidis JPA. Global perspective of COVID-19 epidemiology for a full-cycle pandemic. Eur J Clin Invest. 2020:e13421. Epub 2020/10/08. doi: 10.1111/eci.13423. PubMed PMID: 33026101.

39. Nazarullah A, Liang C, Villarreal A, Higgins RA, Mais DD. Peripheral Blood Examination Findings in SARS-CoV-2 Infection. Am J Clin Pathol. 2020;154(3):319–29. Epub 2020/08/07. doi: 10.1093/ajcp/aqaa108. PubMed PMID: 32756872; PubMed Central PMCID: PMCPMC7454310.

40. Zheng M, Gao Y, Wang G, Song G, Liu S, Sun D, et al. Functional exhaustion of antiviral lymphocytes in COVID-19 patients. Cellular & Molecular Immunology. 2020;17(5):533–5. doi: 10.1038/s41423-020-0402-2.

41. Channappanavar R, Fehr Anthony R, Vijay R, Mack M, Zhao J, Meyerholz David K, et al. Dysregulated Type I Interferon and Inflammatory Monocyte-Macrophage Responses Cause Lethal Pneumonia in SARS-CoV-Infected Mice. Cell Host & Microbe. 2016;19(2):181–93. doi: https://doi.org/10.1016/j.chom.2016.01.007.

42. Channappanavar R, Fehr AR, Zheng J, Wohlford-Lenane C, Abrahante JE, Mack M, et al. IFN-I response timing relative to virus replication determines MERS coronavirus infection outcomes. The Journal of Clinical Investigation. 2019;129(9):3625–39. doi: 10.1172/JCI126363.

43. Blanco-Melo D, Nilsson-Payant BE, Liu W-C, Uhl S, Hoagland D, Møller R, et al. Imbalanced Host Response to SARS-CoV-2 Drives Development of COVID-19. Cell. 2020;181(5):1036–45.e9. doi: 10.1016/j.cell.2020.04.026.

44. Fan BE, Chong VCL, Chan SSW, Lim GH, Lim KGE, Tan GB, et al. Hematologic parameters in patients with COVID-19 infection. American Journal of Hematology. 2020;95(6):E131–E4. doi: 10.1002/ajh.25774.

45. Taneri PE, Gómez-Ochoa SA, Llanaj E, Raguindin PF, Rojas LZ, Roa-Díaz ZM, et al. Anemia and iron metabolism in COVID-19: a systematic review and meta-analysis. European Journal of Epidemiology. 2020;35(8):763–73. doi: 10.1007/s10654-020-00678-5.

46. Richardson S, Hirsch JS, Narasimhan M, Crawford JM, McGinn T, Davidson KW, et al. Presenting Characteristics, Comorbidities, and Outcomes Among 5700 Patients Hospitalized With COVID-19 in the New York City Area. JAMA. 2020;323(20):2052–9. doi: 10.1001/jama.2020.6775.

47. Kernan KF, Carcillo JA. Hyperferritinemia and inflammation. Int Immunol. 2017;29(9):401–9. Epub 2017/05/26. doi: 10.1093/intimm/dxx031. PubMed PMID: 28541437; PubMed Central PMCID: PMCPMC5890889.

48. Wessling-Resnick M. Crossing the Iron Gate: Why and How Transferrin Receptors Mediate Viral Entry. Annu Rev Nutr. 2018;38:431–58. Epub 2018/06/01. doi: 10.1146/annurev-nutr-082117-051749. PubMed PMID: 29852086; PubMed Central PMCID: PMCPMC6743070.

49. Cassat James E, Skaar Eric P. Iron in Infection and Immunity. Cell Host & Microbe. 2013;13(5):509–19. doi: 10.1016/j.chom.2013.04.010.

50. Matsuyama S, Nagata N, Shirato K, Kawase M, Takeda M, Taguchi F. Efficient activation of the severe acute respiratory syndrome coronavirus spike protein by the transmembrane protease TMPRSS2. J Virol. 2010;84(24):12658–64. Epub 2010/10/12. doi: 10.1128/JVI.01542-10. PubMed PMID: 20926566; PubMed Central PMCID: PMCPMC3004351.

51. Mason RJ. Pathogenesis of COVID-19 from a cell biology perspective. European Respiratory Journal. 2020;55(4):2000607. doi: 10.1183/13993003.00607-2020.

52. Nakajima N, Asahi-Ozaki Y, Nagata N, Sato Y, Dizon F, Paladin FJ, et al. SARS coronavirus-infected cells in lung detected by new in situ hybridization technique. Jpn J Infect Dis. 2003;56(3):139–41. Epub 2003/08/29. PubMed PMID: 12944688.

53. Martines R, Ritter J, Matkovic E, Gary J, Bollweg B, Bullock H, et al. Pathology and Pathogenesis of SARS-CoV-2 Associated with Fatal Coronavirus Disease, United States. Emerging Infectious Disease journal. 2020;26(9). doi: 10.3201/eid2609.202095.

54. Ledford H. Coronavirus reinfections: three questions scientists are asking. Nature. 2020;585(7824):168–9. Epub 2020/09/06. doi: 10.1038/d41586-020-02506-y. PubMed PMID: 32887957.

55. Chandrashekar A, Liu J, Martinot AJ, McMahan K, Mercado NB, Peter L, et al. SARS-CoV-2 infection protects against rechallenge in rhesus macaques. Science. 2020. Epub 2020/05/22. doi: 10.1126/science.abc4776. PubMed PMID: 32434946; PubMed Central PMCID: PMCPMC7243369.

56. Seow J, Graham C, Merrick B, Acors S, Pickering S, Steel KJA, et al. Longitudinal observation and decline of neutralizing antibody responses in the three months following SARS-CoV-2 infection in humans. Nature Microbiology. 2020. doi: 10.1038/s41564-020-00813-8.

57. Long QX, Liu BZ, Deng HJ, Wu GC, Deng K, Chen YK, et al. Antibody responses to SARS-CoV-2 in patients with COVID-19. Nat Med. 2020;26(6):845–8. Epub 2020/05/01. doi: 10.1038/s41591-020-0897-1. PubMed PMID: 32350462.

58. Long QX, Tang XJ, Shi QL, Li Q, Deng HJ, Yuan J, et al. Clinical and immunological assessment of asymptomatic SARS-CoV-2 infections. Nat Med. 2020. Epub 2020/06/20. doi: 10.1038/s41591-020-0965-6. PubMed PMID: 32555424.

59. Sekizuka T, Itokawa K, Hashino M, Kawano-Sugaya T, Tanaka R, Yatsu K, et al. A genome epidemiological study of SARS-CoV-2 introduction into Japan. medRxiv. 2020:2020.07.01.20143958. doi: 10.1101/2020.07.01.20143958.

60. Koyama T, Platt D, Parida L. Variant analysis of SARS-CoV-2 genomes. Bull World Health Organ. 2020;98(7):495–504. Epub 2020/08/04. doi: 10.2471/BLT.20.253591. PubMed PMID: 32742035; PubMed Central PMCID: PMCPMC7375210.

61. Rouchka EC, Chariker JH, Chung D. Variant analysis of 1,040 SARS-CoV-2 genomes. PLoS One. 2020;15(11):e0241535. Epub 2020/11/06. doi: 10.1371/journal.pone.0241535. PubMed PMID: 33152019; PubMed Central PMCID: PMCPMC7643988.

62. Matsuyama S, Nao N, Shirato K, Kawase M, Saito S, Takayama I, et al. Enhanced isolation of SARS-CoV-2 by TMPRSS2-expressing cells. Proc Natl Acad Sci U S A. 2020;117(13):7001–3. Epub 2020/03/14. doi: 10.1073/pnas.2002589117. PubMed PMID: 32165541; PubMed Central PMCID: PMCPMC7132130.

63. Itokawa K, Sekizuka T, Hashino M, Tanaka R, Kuroda M. Disentangling primer interactions improves SARS-CoV-2 genome sequencing by multiplex tiling PCR. PLoS One. 2020;15(9):e0239403. Epub 2020/09/19. doi: 10.1371/journal.pone.0239403. PubMed PMID: 32946527; PubMed Central PMCID: PMCPMC7500614.

64. Li H, Durbin R. Fast and accurate short read alignment with Burrows-Wheeler transform. Bioinformatics. 2009;25(14):1754–60. Epub 2009/05/20. doi: 10.1093/bioinformatics/btp324. PubMed PMID: 19451168; PubMed Central PMCID: PMCPMC2705234.

65. Coil D, Jospin G, Darling AE. A5-miseq: an updated pipeline to assemble microbial genomes from Illumina MiSeq data. Bioinformatics. 2015;31(4):587–9. Epub 2014/10/24. doi: 10.1093/bioinformatics/btu661. PubMed PMID: 25338718.

66. Shirato K, Nao N, Katano H, Takayama I, Saito S, Kato F, et al. Development of Genetic Diagnostic Methods for Novel Coronavirus 2019 (nCoV-2019) in Japan. Jpn J Infect Dis. 2020. Epub 2020/02/20. doi: 10.7883/yoken.JJID.2020.061. PubMed PMID: 32074516.

67. Babraham AS. Bioinformatics - FastQC A Quality Control tool for High Throughput Sequence Data [cited 24 Sep 2020]. Available from: https://www.bioinformatics.babraham.ac.uk/projects/fastqc/.

68. Bolger AM, Lohse M, Usadel B. Trimmomatic: a flexible trimmer for Illumina sequence data. Bioinformatics. 2014;30(15):2114–20. Epub 2014/04/04. doi: 10.1093/bioinformatics/btu170. PubMed PMID: 24695404; PubMed Central PMCID: PMCPMC4103590.

69. Dobin A, Davis CA, Schlesinger F, Drenkow J, Zaleski C, Jha S, et al. STAR: ultrafast universal RNA-seq aligner. Bioinformatics. 2013;29(1):15–21. Epub 2012/10/30. doi: 10.1093/bioinformatics/bts635. PubMed PMID: 23104886; PubMed Central PMCID: PMCPMC3530905.

70. Liao Y, Smyth GK, Shi W. featureCounts: an efficient general purpose program for assigning sequence reads to genomic features. Bioinformatics. 2014;30(7):923–30. Epub 2013/11/15. doi: 10.1093/bioinformatics/btt656. PubMed PMID: 24227677.

71. Anders S, Huber W. Differential expression analysis for sequence count data. Genome Biol. 2010;11(10):R106. Epub 2010/10/29. doi: 10.1186/gb-2010-11-10-r106. PubMed PMID: 20979621; PubMed Central PMCID: PMCPMC3218662.

72. CRAN. Package pheatmap [cited 24 Sep 2020]. Available from: https://cran.r-project.org/web/packages/pheatmap/index.html.

73. Weiner 3rd J DT. tmod: an R package for general and multivariate enrichment analysis. PeerJ Preprints. 2016. doi: 10.7287/peerj.preprints.2420v1.

74. Smyth GK. limma: Linear Models for Microarray Data. In: Gentleman R. CVJ, Huber W., Irizarry R.A., Dudoit S., editor. Bioinformatics and Computational Biology Solutions Using R and Bioconductor Springer, New York, NY: Springer, New York, NY; 2005. p. 397–420.

